# Causal variant capture in genotype discovery approaches drives polygenic prediction performance across traits and populations

**DOI:** 10.1101/2025.03.18.644029

**Authors:** Yi-Sian Lin, Taotao Tan, Ying Wang, Bogdan Pasaniuc, Alicia R. Martin, Elizabeth G. Atkinson

**Author notes:** **Corresponding author contact information:** Baylor College of Medicine, One Baylor Plaza, Taub Research Building Room 621, Houston, TX, USA 77030.

## Abstract

Polygenic scores (PGS) are widely used to estimate genetic predisposition to complex traits by aggregating the effects of common variants into a single measure. They hold promise in identifying individuals at increased risk for diseases, allowing earlier screening and interventions. Genotyping arrays, commonly used for PGS computation, are affordable and computationally efficient, while whole-genome sequencing (WGS) offers a more comprehensive view of genetic variation. In this study, we compared PGS derived from arrays and WGS across multiple traits to evaluate differences in predictive performance, portability across populations, and computational efficiency. We computed PGS for 10 traits, representing a range of heritability and polygenicity, in the three largest genetic ancestry groups in *All of Us* (European, African American, Admixed American), trained on multi-ancestry meta-analyses from the Pan-UK Biobank. Using the clumping and thresholding (C+T) method, we found that WGS-based PGS outperformed array-based PRS for highly polygenic traits but showed differentially reduced accuracy for sparse traits in certain populations. With the LD-informed PRS-CS method, we observed overall improved prediction performance compared to C+T, with WGS outperforming arrays across most non-cancer traits. The results obtained using PRS-CS closely align with those derived from pre-trained models in the PGS Catalog, with prediction achieving better performance using WGS than array genotypes for non-sparse traits. To further investigate factors influencing differential prediction performance between array and WGS, we ran simulations varying the proportions of causal SNPs directly captured by the technologies. These demonstrated that the proportion of causal variants genotyped dramatically affects prediction accuracy. Fine-mapping of empirical data supported this concept but also highlighted the importance of reducing non-informative variants for optimal prediction accuracy. In conclusion, while WGS-based PGS generally offer superior predictive power with PRS-CS, the advantage over arrays is context-dependent, varying by trait, population, and the PGS method. The ability to capture causal variants through these technologies largely drives the prediction accuracy. This study provides insights into the complexities and potential advantages of using different genotype discovery approaches for polygenic predictions across populations and informs on strategies to enhance accuracy.

## Introduction

Polygenic scores (PGS), also known as polygenic risk scores (PRS), estimate an individual’s genetic predisposition to complex traits or diseases. As an emerging tool in precision medicine, PGS holds significant potential for informing early disease detection, risk stratification, prevention, and intervention^1^. PGS are calculated by aggregating the genotypes (i.e., the dosage of minor or risk alleles) from a target cohort, weighted by their corresponding effect sizes derived from relevant genome-wide association studies (GWAS) conducted in a discovery cohort^2^. The prediction accuracy of PGS is influenced by several factors, such as the sample size of the discovery cohort and the genetic similarity between the discovery and target population^3,4^. However, with up to 90% of GWAS for complex traits conducted in populations with predominantly European ancestry^5,6^, PGS estimates are often less accurate for individuals of non-European ancestries, which can contribute to health disparities across populations^7^.

Genotype data from both discovery and target cohorts are typically obtained using array technologies that detect known genetic polymorphisms. Since 2005, Affymetrix has developed high-throughput microarray genotyping platforms widely used in early GWAS and PGS studies^8,9^, which initially assayed between 500,000 and nearly a million SNPs^10^. More recently, Illumina’s Omni series arrays expanded this coverage to detect millions of genetic markers^11^. In addition to capturing genetic variation in European populations, arrays designed for global and diverse populations, such as the Global Screening Array^12^, Global Diversity Array^13,14^, Asian Screening Array^15^, and H3 Africa Array^16,17^, became available following the completion of the 1000 Genomes Project (TGP) Phase 3^18^ and other diverse reference resources. To allow for more comprehensive genome-wide coverage, imputation is often employed to infer untyped variants, thereby enhancing the accuracy of GWAS and PGS predictions^19^.

With the ongoing reduction in costs and advancements in high-throughput sequencing, both whole exome sequencing (WES) and whole genome sequencing (WGS) have become increasingly integrated into population-level genomic studies and clinical applications^20,21^. In contrast to array-based genotyping, sequencing offers several distinct advantages. Unlike genotyping arrays, which are typically designed to assay common variants and require imputation to enhance coverage, deep sequencing captures all forms of genetic variants ranging from ultra-rare to common alleles^22,23^. Additionally, the imputation accuracy for genotyping arrays can impact the predictive performance of PGS^24^, particularly in non-European populations. This is due to their inability to reliably impute variants with low frequency, and variants enriched in populations that are not well represented in existing reference panels. WGS enables the direct assessment of low-frequency variants and loci that are missed or poorly imputed in arrays. Therefore, these advantages could potentially lead to more accurate and robust PGS, especially for underrepresented populations and traits influenced by rarer variants. However, WGS remains more expensive, and generates vast amounts of data which requires computationally intensive processing.

Previous studies have demonstrated comparable results between low-coverage sequencing and array-based genotyping in gene discovery^25–27^ and PGS for diseases like Parkinson’s disease^28^. However, few studies have assessed the impact of different genotype discovery technologies on PGS performance at the biobank scale or considered high-coverage WGS versus arrays. The *All of Us* Research Program^14^ (v6), with 95,562 samples with both array and WGS data, provides a unique opportunity to evaluate the predictive accuracy across technologies in multiple ancestral groups and numerous phenotypes. In this study, we benchmarked the performance of PGS derived from genotyping arrays, pre-curated models released in the widely-used PGS catalog, and 30x high-coverage WGS across up to ten traits and multiple populations in *All of Us* (Fig. 1a). Through these efforts, we aim to assess the conditions under which WGS-based PGS outperforms array-based PGS and identify trade-offs that may inform the choice of genomic data type for risk prediction. We also test the hypothesis that differences in the proportion of causal variants captured explains the observed differences in prediction accuracy across technologies, providing insights into improving polygenic prediction (Fig. 1b).

**Fig. 1:**
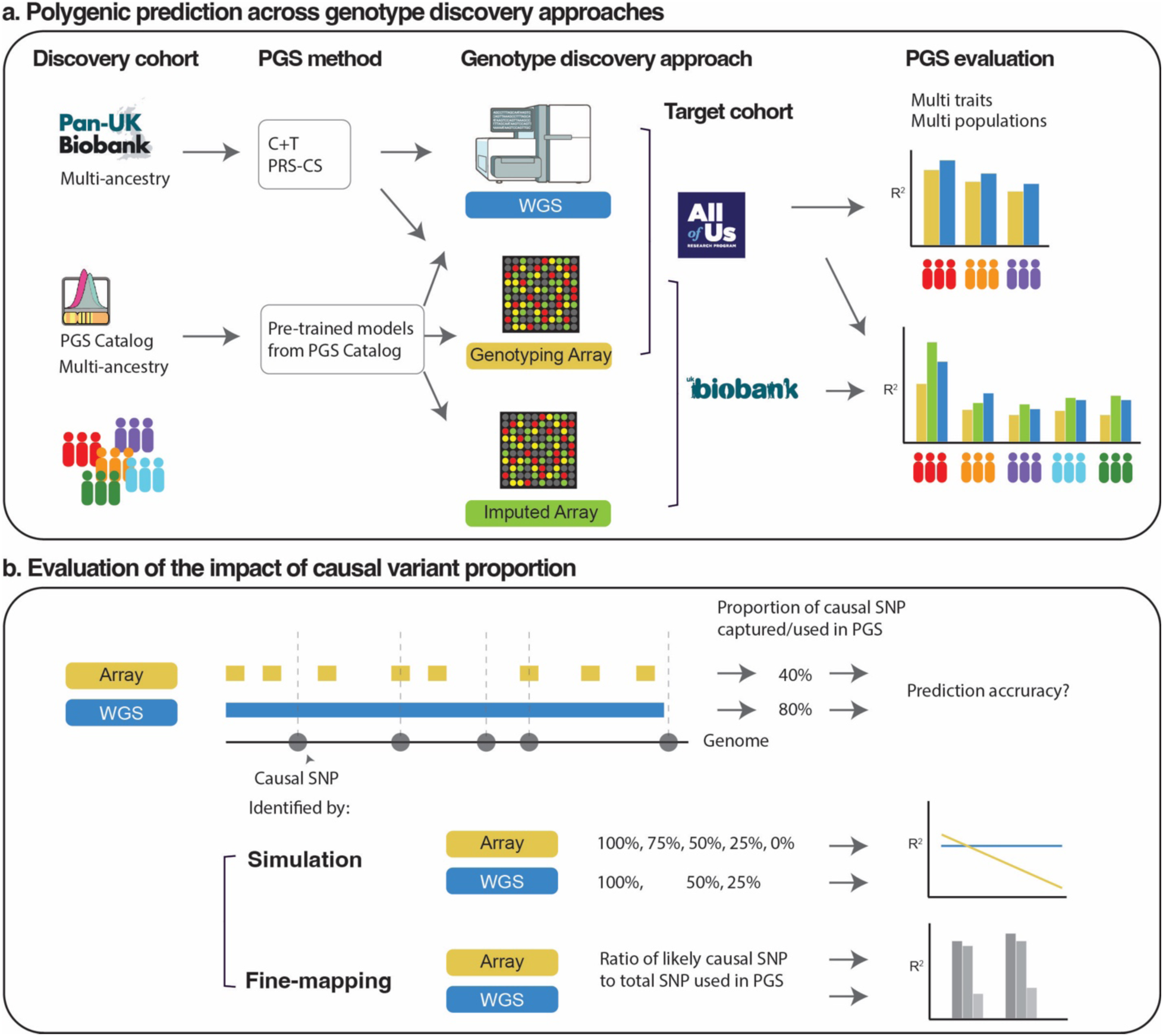
Study design. In the phase (a), PGS were calculated for each *All of Us* individual in the EUR, AMR, or AFR populations using genotypes derived from either array or WGS approaches. C+T and PRS-CS methods were benchmarked. Pre-trained models from the PGS Catalog were used to evaluate performance in *All of Us* and UKBB individuals across multiple populations, using genotypes derived from direct genotyping, imputation, or WGS. In the phase (b), prediction performance was evaluated based on different proportions of causal variants captured by the genotype discovery approach, from simulation and fine-mapping.

## Results

### Selection of traits across a range of genetic architectures

The primary goal of this study is to benchmark the predictive performance from array-based versus sequencing-based genotypes. We first selected 95,562 target samples from the *All of Us* v6 dataset, each with both array and WGS genotype data. PGS are expected to exhibit higher predictive accuracy when the ancestry of the discovery and target cohorts is matched^29^. However, since this study aims to develop a generalized PGS model rather than focusing on a single ancestry, we used effect sizes derived from multi-ancestry GWAS meta-analyses of the Pan-UKBB cohort. For the target samples in *All of Us*, we focused on European (EUR), Latino/Admixed American (AMR), and African/African American (AFR) cohorts based on predicted genetic ancestry (Supplementary Fig. 1), as these groups had sample sizes sufficiently large to support multi-trait comparative analyses. East Asian (EAS) and South Asian (SAS) populations were included in some selected trait analyses if the sample size was sufficient.

We selected 10 complex traits with distinct genetic architectures (Supplementary Table 1) from the Pan-UKBB and their equivalents in the *All of Us* dataset (Supplementary Table 2): standing height (Height), diastolic blood pressure (DBP), white blood cell count (Leukocyte), red blood cell count (RBC), high-density lipoprotein cholesterol (HDL), total cholesterol (TC), asthma, breast cancer, colorectal cancer, and type 2 diabetes (T2D). Height exhibits the highest heritability and polygenicity among the traits, followed by HDL, whereas TC and asthma are the least heritable, and cancers are the sparsest traits (Supplementary Fig. 2, Supplementary Table 3).

### WGS improves prediction over arrays with LD-informed effect size shrinkage

We first assessed the PGS using the classic clumping and thresholding (C+T) method (Supplementary Table 4 and 5). Notably, the raw average PGS for array-based and WGS-based genotypes were highly correlated across traits (*r* > 0.9 for all traits except for height and TC), suggesting that both platforms captured similar genetic information, resulting in comparable polygenic predictions. This also indicates that the genotype coverage from unimputed array data is sufficient for constructing reliable PGS comparable to those from WGS in the *All of Us* dataset for most of the traits considered. Furthermore, it is noteworthy that EUR populations tend to show smaller and more homogenous raw PGS, whereas AFR populations tend to exhibit larger and more widely spread PGS, particularly for colorectal cancer and breast cancer (Supplemental Fig. 3).

The predictive accuracy, measured by R^2^ or Nagelkerke’s R^2^, was assessed across different genotype discovery approaches and populations (Fig. 2, Supplementary Table 6 and 7). Given that EUR individuals constitute a large proportion of both the discovery cohort (95%) and the target cohort (52%), predictive accuracy in EUR was generally higher than in other populations, as expected, regardless of the genotype discovery technology (ranging from 1.58 × 10^−3^ ± 1.28 × 10^−3^ to 1.07 × 10^−1^ ± 6.12 × 10^−4^). AFR populations generally showed lower performance compared to other groups (ranging from 7.33 × 10^−4^ ± 7.53 × 10^−4^ to 3.37 × 10^−2^ ± 1.07 × 10^−3^, Supplementary Table 8). Surprisingly, PGS performance from WGS genotypes was suboptimal relative to array genotypes for multiple traits, and only outperformed arrays in all three populations for one trait - height. This discrepancy may be due to height’s highly polygenic nature, suggesting that the higher number of variants detected by WGS provides more informative variants for PGS prediction than array genotypes. Additionally, there are a few phenotypes with notable differences in R² across genotype discovery approaches. In addition to height, prediction from WGS also performed better for TC, RBC, and leukocyte count in EUR, and for leukocyte count and asthma in AFR. In contrast, we observed that PGS derived from array genotypes performed better for several other traits, particularly HDL in AMR (average array-based R^2^=0.061; average WGS-based R^2^=0.048; paired t-test *p* = 3.43 × 10⁻⁴) and AFR populations (average array-based R^2^=0.027; average WGS-based R^2^=0.009; paired t-test *p* = 7.7 × 10⁻⁷), as well as RBC in AFR, T2D in AMR, and breast cancer in EUR (Supplementary Table 8 and 9). Several findings from previous studies may provide insights for interpreting these observations, including aspects of population-enriched architecture, as detailed in Supplementary Table 10.

**Fig. 2:**
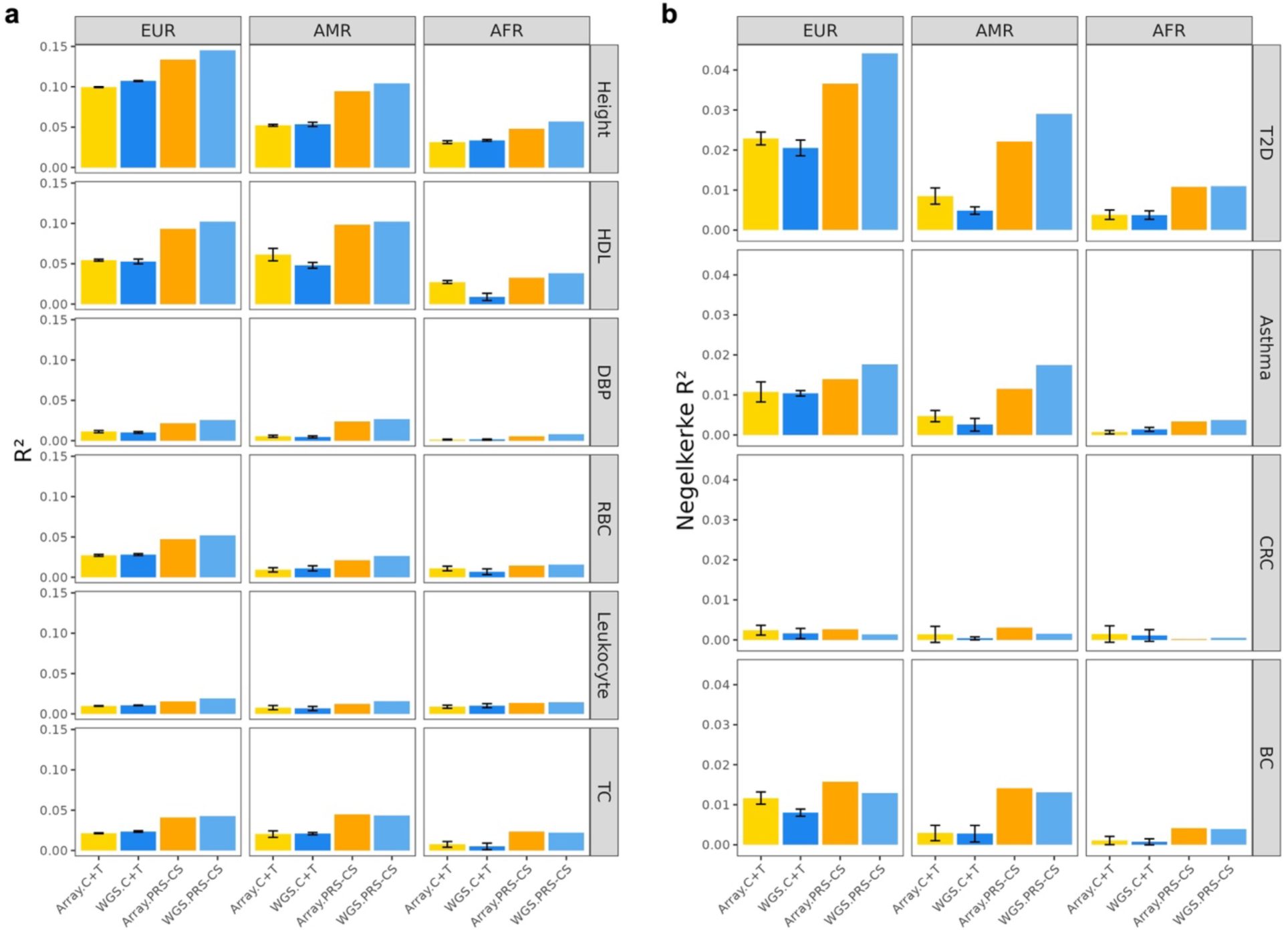
Comparison of PGS performance across genotype discovery approaches. PGS were computed for samples with genotypes derived from arrays (yellow, orange) or WGS (blues) in *All of Us*. Continuous traits (a) and binary traits (b) are ordered by polygenicity. R² or Nagelkerke’s R² were determined by adjusting for age, sex, and the first 16 genetic PCs. Black bars indicate 95% confidence intervals. HDL: high-density lipoprotein cholesterol. DBP: diastolic blood pressure. RBC: red blood cell count. TC: total cholesterol. T2D: type 2 diabetes. CRC: colorectal cancer. BC: breast cancer.

C+T retains only quasi-independent variants within each LD block after LD clumping; however, this approach may remove informative variants important for PGS calculation. In contrast, PRS-CS does not rely on clumping but instead continuously shrinks effect size estimates in an LD-informed manner to retain as many informative variants as possible^30^. We benchmarked C+T and PRS-CS to determine whether prediction accuracy would differ between the two methods across genotype discovery technologies. For raw PGS using PRS-CS, we again observed a high correlation between PGS derived from array-based genotypes and WGS genotypes across traits (*r* > 0.9 for continuous traits) (Supplementary Fig. 4). When comparing the predictive accuracy of PGS generated using PRS-CS to those generated using the C+T method, we found that PRS-CS generally outperformed C+T, with performance being approximately two times better for most traits, except for cancers. Additionally, performance using WGS variants was higher than that using array variants for all traits except for cancers and TC (Fig. 2).

### HapMap3 variants are sufficient for PRS-CS prediction compared with full WGS panels

Note that the variants selected for PGS computation were restricted to approximately 1.1 million HapMap3 variants, as the default LD panel used in PRS-CS was based on HapMap3 variants. This is reflective of the general implementation of the PRS-CS software. To fully assess the advantage of using WGS genotypes, however, which include around 9 million variants, we additionally computed an EUR-derived LD matrix using samples from the Thousand Genomes Project (TGP) and the Human Genome Diversity Project (HGDP) joint call sets, resulting in approximately 7.3 million reference variants. (Supplementary Table 11). Switching from the default LD panel to this new panel, we observed similar levels of predictive accuracies for traits using array genotypes, but, surprisingly, lower accuracy using WGS variants in AMR and AFR for multiple traits (Supplementary Fig. 5, Supplementary Table 12). This could be attributable to the higher number of variants in the EUR LD panel, which does not tag well for non-EUR population. However, taken together, LD information from HapMap3 variants is generally sufficient to perform PRS-CS and achieve optimal performance without the computationally demanding need to construct fuller LD resources.

### Pre-trained PGS show performance gains from imputation and WGS

PGS methods can substantially influence predictive accuracy. We next evaluated the performance of pre-trained PGS models on genotypes called from different technologies. Specifically, we computed PGS using (posterior) effect size estimates reported in PGS Catalog^31^ studies for four traits: height^32,33^, HDL^34,35^, T2D^36^, and breast cancer^37^. This analysis enabled assessment of the generalizability of prediction performance across genotype discovery approaches using a third PGS framework (Supplementary Table 13). In addition to evaluation in the *AoU* cohort, we applied the same models to the UK Biobank (UKBB) cohort, allowing comparison of array-based predictions both before and after genotype imputation.

Several patterns emerge. First, within the same discovery studies, prediction derived from WGS generally outperformed those from array in the *AoU* cohort across most all traits and populations, with exception of breast cancer in AMR, EAS, and SAS populations. Similarly, in UKBB, imputed array genotypes consistently yielded higher predicative accuracy than non-imputed arrays across all traits and populations (Figure 3 hashed bars). Second, the performance gain from non-imputed to imputed array in UKBB was generally larger than the gain observed from array to WGS in *AoU*. Although WGS is expected to provide a less biased and more accurate representation of genetic variation, the apparent advantage of imputation may reflect the introduction of dense LD structure that inflate prediction, and these results should therefore be interpretated with caution. Third, we compared height PGS from two independent studies. Scores from Yengo *et al.* showed higher predictive performance in UKBB than in *AoU*, whereas scores from Gunn *et al.* performed better in *AoU* than in UKBB. Notably, the original PGS from Yengo *et al.* was evaluated in UKBB, while the PGS from Gunn *et al.* was originally evaluated in *AoU*. This demonstrates that prediction performance is higher in the cohort where tuning occurred.

**Fig. 3:**
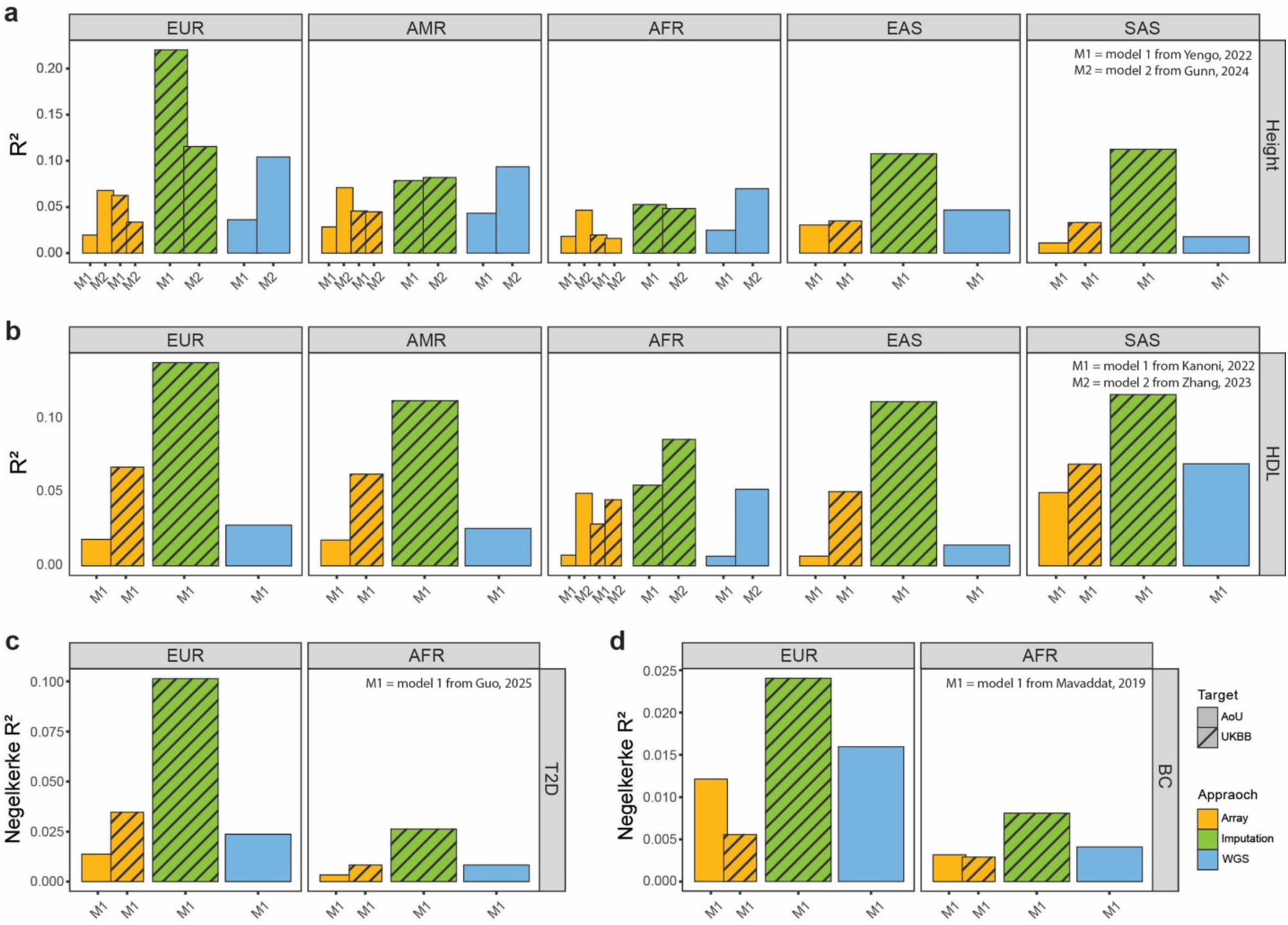
Evaluation of PGS performance from the PGS Catalog. Polygenic score data for four traits, (a) height, (b) HDL, (c) T2D, and (d) breast cancer, were extracted from the PGS Catalog. The studies for each trait, where PGS models were trained, are labeled in the upper right corner. Performance was evaluated using genotype discovery approaches from two sources: array and WGS data from *AoU*, and direct genotyping array and imputation data from UKBB. The central South Asian (CSA) group in UKBB was renamed to the SAS group for comparison with *AoU*. For T2D and breast cancer, results are shown only for EUR and AFR populations due to insufficient sample sizes (n < 100) in AMR within UKBB and in EAS and SAS within *AoU*. BC: breast cancer.

### Causal variant capture shapes polygenic prediction

We next investigated factors influencing differential predictive performance across genotype discovery approaches. We hypothesized that prediction accuracy increases with the proportion of causal variants captured by a given genotype discovery technology (Fig. 1b). To test this hypothesis, we simulated polygenic scenarios in which array and WGS data captured different proportions of causal variants (total number of causal variants M = 500, ∼0.1% of the studied genomic variants, Fig. 4a). Using *AoU* genotypes for simulation, we observed a decrease in prediction R^2^ as the proportion of captured causal variants decreased for both WGS (100%, 50%, or 25%) and array data (100%, 75%, 50%, 25%, or 0%). This dose-response pattern remained consistent under a less polygenic setting (M = 50, Fig. 4b). Surprisingly, when using UKBB genotypes for simulation, reducing the proportion of causal variants used by imputed array did not result in a corresponding decrease in predictive performance (Supplementary Fig. 6). This discrepancy may reflect inflated performance due to introduction of dense LD structure during imputation, consistent with observations from real data analyses (Fig. 3). Overall, these results indicate that the proportion of causal variants captured is a key determinant of polygenic prediction accuracy.

**Fig. 4:**
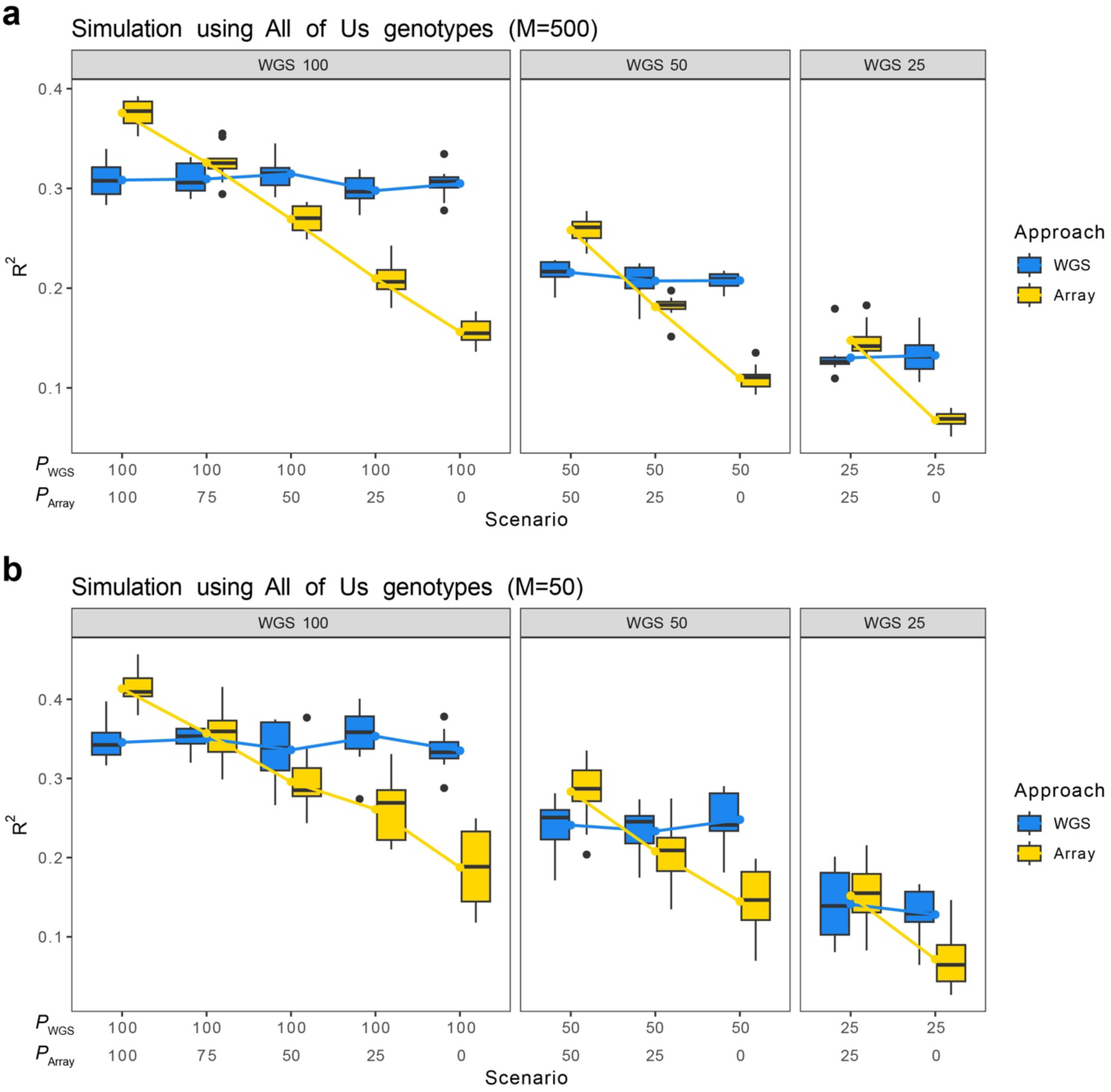
Simulation settings of causal variants captured. Traits were simulated using *AoU* WGS genotypes under polygenic settings (**a**, with *M* = 500 causal variants) and sparse settings (**b**, with *M* = 50 causal variants), both with heritability = 0.5. Ten scenarios were simulated. *P*_WGS_ represent the percentage of causal variants captured by WGS, and *P*_Array_ denote the percentage of causal variants captured by array.

To test this concept in real data, we performed statistical fine-mapping using SuSiE to infer likely causal variants for four traits - height, HDL, T2D, and breast cancer – using the Pan-UKBB summary statistics for EUR. In total, we identified 1,185 likely causal SNPs for height, 225 for HDL, 8 for T2D, and 3 for breast cancer (Supplementary Table 14-17). We then quantified the proportion of those inferred causal SNPs included in the default PRS-CS framework (restricted to HapMap3 variants) and in PRS-CS using the extended LD reference panel. For T2D, the proportion of inferred causal variants captured by array increased from 12% using the HapMap3 SNP set to 38% using the full variant set, with no noticeable increase for other traits (Supplementary Fig. 7a). In contrast, for WGS data, the proportion of inferred causal SNPs used increased substantially when extending the variant set beyond HapMap3, indicating that WGS captures a large number of likely causal variants outside the HapMap3 subset (Supplementary Fig. 7b). However, capturing a higher proportion of causal variants alone did not translate into improved prediction accuracy (Fig. 5), potentially due to the inclusion of additional noise from uninformative SNPs in the model. These results indicate that the signal-to-noise ratio, rather than the absolute number of causal variants captured, might be a key determinant of relative performance between array- and WGS-based PGS. To further investigate this hypothesis, we ran PRS-CS using only inferred causal variants while excluding all other variants. Unexpectedly, prediction performance decreased substantially, suggesting that either many true causal variants are missed by statistical fine-mapping or that additional or more common variants correlated with casual SNPs (for example, through LD) are required for improved polygenic prediction.

**Fig. 5:**
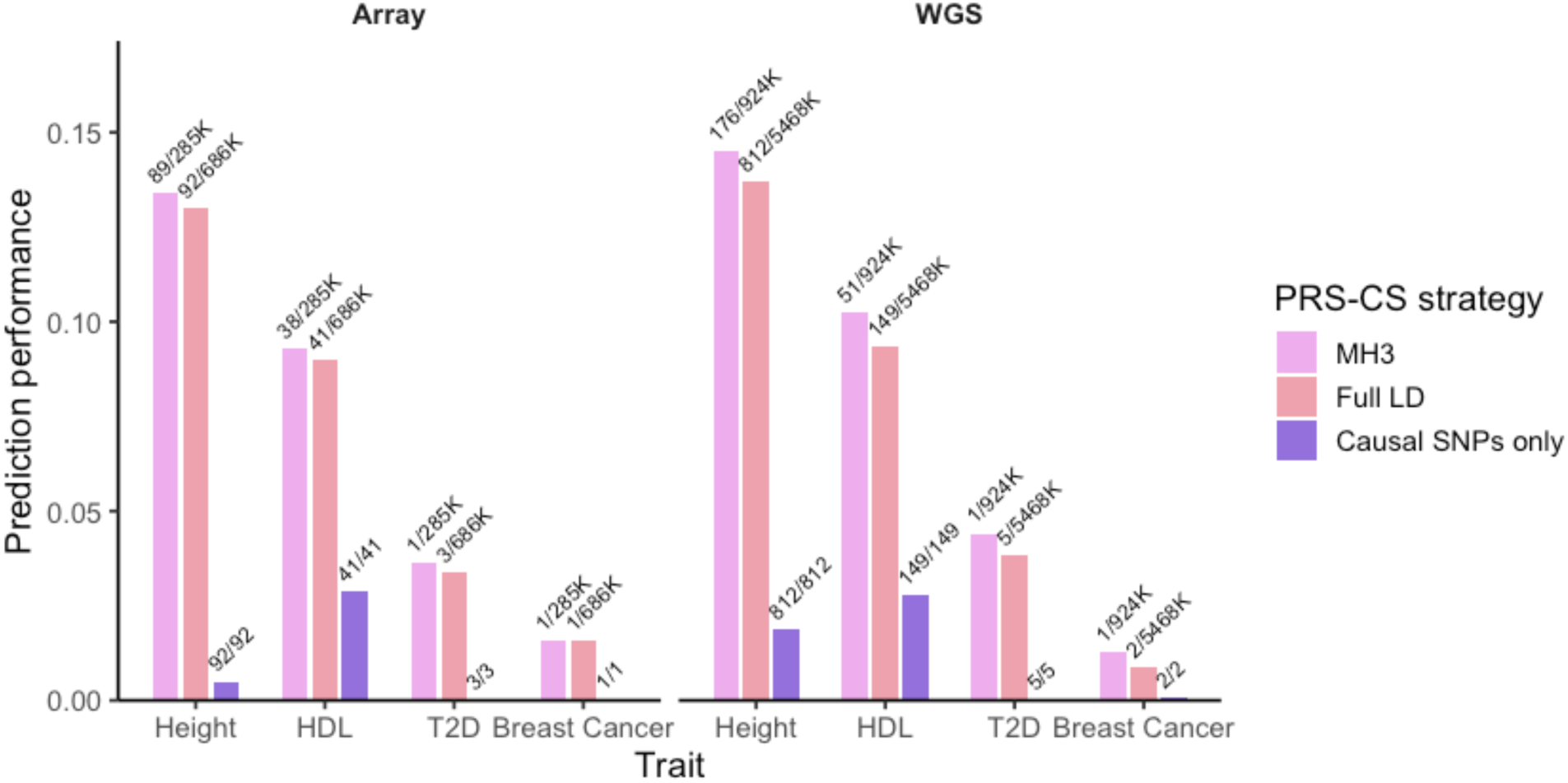
Prediction performance of PGS including different numbers of causal variants. The performance for selected traits was assessed with three PRS-CS strategies for variant selection: (HM3) variants intersected with the LD panel constructed using HapMap3 variants (default in PRS-CS), (Full LD) variants intersected with a full LD reference panel, and (Causal SNPs only) only inferred causal SNPs were used with the full LD panel. The numbers on top of each bar indicate the number of causal variants used, out of (/) the total variants included in the PRS-CS construction.

### Trade-off between computational efficiency and prediction accuracy

From a practical perspective, the choice of genotype discovery approach involves important trade-offs between predictive performance, data generation cost, and computational burden. The cost of genotyping arrays is around $100 per sample, whereas 30X WGS is about 5-6 times more expensive. Computationally, PGS construction using array genotypes was much more efficient than using WGS data. Focusing on post-QC computation with C+T, WGS required approximately twice the CPU time per trait compared with array data in the *AoU* dataset (∼54 vs. ∼27 CPU hours) for LD clumping and selection of the optimal PGS via 10-fold cross-validation. For PRS-CS, inferring posterior effect sizes using WGS required 2.2-fold more CPU time than array (175.5 vs. 81 CPU hours) on a local server outside of *AoU*, and up to an 18-fold increase in CPU time using a full-sized LD matrix (WGS: 5,616 CPU hours; array: 306 CPU hours; Supplementary Table 18). Subsequent PGS computation within *AoU* environment was also more computationally expensive for WGS than for array data. Together, these results highlight data generation and computational cost as key considerations when selecting a genotype discovery approach for PGS analyses.

In conclusion, polygenic prediction performance varies across genotype discovery approaches, PGS methods, traits, and populations. WGS data, when used with LD-informed methods such as PRS-CS, generally yields higher predictive power than array data, particularly for traits with higher heritability. However, array-derived PGS remain more cost-effective and computationally efficient, making them a practical choice for rapid PGS generation and more accurate for traits with sparser genetic architectures, such as cancers. Consistent with our primary analyses, evaluation using pre-trained models from the PGS Catalog recapitulate similar performance patterns across genotype discovery technologies, while providing a computationally efficient alternative for rapid PGS assessment. However, the accuracy of pre-trained models is generally lower than that of self-trained models. Finally, our results underscore that capturing causal variants is critical for polygenic prediction, whereas the inclusion of noninformative variants negatively influence performance across genotype discovery approaches.

## Discussion

In this study, we systematically benchmarked the prediction accuracy of PGS generated from array and WGS genotypes across multiple traits and populations in the *All of Us* cohort. We show that the relative performance of these approaches depends on the interplay between genotype discovery technology, PGS method, trait architecture, and ancestry. By comparing our results, particularly PRS-CS (Fig. 2), with pre-trained scores from the PGS Catalog, we demonstrate the consistency that WGS-based PGS generally outperforms array-based scores for non-sparse traits when using Bayesian methods. This suggests that these results are generalize across more advanced modeling strategies. Furthermore, through direct comparison with the paired UKBB array data before and after imputation array, we illustrate that imputation can achieve or even exceed the predictive gains of WGS relative to unimputed arrays (Fig. 3).

Using the C+T method, WGS generally did not outperform arrays. (Fig. 2). This could potentially be due to the LD clumping, which drastically reduced the number of WGS variants utilized from approximately 9 million to ∼467K (∼94.5% reduction), while array variants clumped down from ∼1 million to ∼262K (a more modest ∼71% reduction) (Supplementary Table 5). The large reduction in WGS variants may have removed important variants for polygenic prediction, leading to comparable or even lower prediction accuracy than PGS derived from array variants. Given that sequencing can better capture additional important genetic variations such as INDELs and CNVs, incorporating more complex genetic variation could boost PGS performance for WGS and recover some missing heritability. Finally, sequencing error rates range from 0.1% to 1% across platforms^38,39^. However, even low error rates can introduce significant errors when large numbers of nucleotides are sequenced^40^. These errors may contribute noise to the PGS, thereby reducing predictive accuracy.

Using PRS-CS, however, WGS-based PGS generally performed better than array-based PGS (Fig. 2). Interestingly, computing our own full LD matrix (rather than utilizing the pre-computed PRS-CS LD matrix that is restricted to HapMap3 SNPs) with 6.2 M additional sites did not improve WGS based results compared to the more limited variant set (Supplementary Fig. 10). These results suggest that fewer, reliably captured, more common, and well-tagging variants appear to suffice for PGS computation for most traits, particularly those that are less polygenic. This can be illustrated with the theoretical accuracy of PGS, where the expected *R*^2^ ≈ *h*^2^⁄[1 + *M*⁄(*h*^2^*N*)]^41^. With the same sample size *N*, using WGS over array variants might increase the SNP heritability *h*^!^, but it does not necessarily provide a benefit when a very large number of variants *M* are involved.

Traditional SNP arrays have predominantly been based on genomes of European ancestry due to the higher availability of EUR samples in reference databases^42^. In contrast, sequencing does not rely on pre-selected markers and is expected to provide greater benefits for underrepresented groups in genomic studies. Consequently, we anticipated a differentially greater improvement in prediction accuracy using variants from sequencing compared to those from arrays in non-European populations. However, this trend was not universally observed, suggesting that the Global Diversity Array used in *All of Us* is not as biased toward European populations as some other arrays are. Ongoing efforts to generate array data for diverse populations would do well to similarly consider capturing variation represented in the target samples.

As has been demonstrated previously^3,43,44^, prediction performance was generally better in EUR than in other populations, likely due to the high representation of EUR in discovery GWAS. Although a low proportion of Pan-UKBB, AMR achieved similar performance to EUR for some traits, while AFR always had low predictive power (Fig. 2). This may be attributed to the larger proportion of European ancestry in the genomes of AMR^45^, and the degree of genetic divergence from EUR, as has been previously explored^4^. We specifically observed a four-fold reduction in performance for T2D in AFR compared to EUR in this study (Fig. 2), highlighting the ongoing performance disparities of PGS for diverse populations across genotype discovery approaches and PGS methods. Another consideration is that AMR and AFR are admixed populations, with genomes composed of genetic segments from multiple ancestries^46^. Ancestry-informed GWAS methods tailored to admixed populations, such as *Tractor* GWAS^47^, may produce summary statistics for polygenic prediction that function better for admixed individuals^29^.

In the PGS Catalog analysis, we observed that imputed genotypes in UKBB confer a huge advantage for prediction performance, particularly in the EUR population (Fig. 3). This is likely because the UKBB imputation relied on reference panels heavily enriched for European samples (e.g., HRC, 1000 Genomes, and UK10K), resulting in more accurate imputation and, consequently, higher prediction accuracy in EUR individuals. Additionally, the imputation process can introduce “artificial” LD, which may inflate apparent predictive gain not only in EUR but in other populations.

In the simulation, we show a clear pattern where a higher proportion of causal variants captured improved prediction performance (Fig. 4). However, in real data, increasing the number of causal variants inferred by statistical fine-mapping did not necessarily lead to better performance, especially when a large number of total variants were included (Fig. 5). When we included only the likely causal SNPs in the PGS, in fact, performance dropped significantly. These findings may suggest that many potential causal variants remain unidentified, and that reducing the number of non-informative variants while increasing informative variants could improve performance.

It is important to note that the current analysis relied on QC’d SNPs, which were common and bi-allelic. Including rare variants and other types of variants could potentially increase the number of likely causal variants identified, provided that appropriate fine-mapping methodologies for these variants are available. The types of variants utilized in both PGS computation and fine-mapping are inherently limited by the variants available in the summary statistics. In our current analysis, the full potential of WGS was restricted by the overlap with summary statistics generated from imputed arrays, which typically represent common variants. If sequencing-based GWAS summary statistics become more prevalent in the future, this is expected to enable broader coverage of variant types in genomic analyses and further improve WGS-based PGS prediction.

The ongoing advancements in sequencing technology may help advance relevant method development. As of 2025, the cost for genotyping arrays ranges from $90 for a screening array to $150 for a global diversity array, while the cost for 30X short-read WGS is about $600. However, the price of WGS has dramatically declined since next-generation sequencing technology came into use in 2007^48^. With the rapid advancement of sequencing technology, WGS is expected to become more affordable and potentially replace arrays as the standard platform in large-scale genomic studies and personal health screening. This development is anticipated to not only accelerate genomic discovery for historically underrepresented populations but also advance PGS method development. The larger and more representative GWAS and reference panels generated from WGS for populations with diverse genetic architectures will enhance prediction accuracy for traits and diseases that have lacked statistical power. This will create many opportunities for genetic prediction models tailored to specific health conditions and populations, facilitating the progress of precision medicine.

To conclude, this study evaluates the performance of PGS using variants identified from different genotype discovery technologies. The advantage of either genotype discovery approach depends on several factors. Array-PGS exhibit higher predictive accuracy than WGS-PGS for sparse traits when variants were clumped. In contrast, WGS and imputation outperformed array-PGS for most traits with higher polygenicity when all variants were used for prediction and effect size estimates were adjusted based on LD. While causal variants drive the prediction power, the inclusion of non-informative variants can negatively impact performance. We hope this study provides researchers with a comprehensive foundation for selecting an optimal genotype calling platform and polygenic scoring method, across diverse populations and trait architectures.

## Methods

### GWAS summary statistics QC

The multi-ancestry meta-analyzed GWAS summary statistics for six continuous traits (height, DBP, leukocyte, RBC, HDL, and TC) and four binary traits (asthma, breast cancer, colorectal cancer, and T2D) were downloaded from Pan-UKBB. In brief, 500,000 UK Biobank samples were genotyped and densely imputed (∼97 million variants)^49^. After filtering for INFO scores > 0.8, approximately 30 million variants were retained and included in the meta-analysis, resulting in a total of 28,987,534 variants in the final summary statistics. A summary table of the selected phenotypes, including sample sizes by population, is provided in Supplementary Table 1. Each summary statistic was lifted over from hg19 to hg38 to harmonize with the genome build of the target genotypes in *All of Us*. Variants in all summary statistics were filtered to retain only biallelic SNPs. After annotating the summary statistics to the Hail table in *All of Us*, mismatches between the summary statistics and target variants were resolved by swapping reference and alternate alleles and/or flipping strands, with corresponding reversal of effect size signs. Ambiguous variants were excluded from further analysis. For quantitative traits, only high-quality samples (denoted with the suffix “hq”), effect sizes, and *P* values were used.

### Heritability and polygenicity estimation

Heritability for the 10 Pan-UKBB phenotypes was estimated using LD Score Regression (LDSC)^50^. Given that EUR constitutes 95% of the total samples in the Pan UKBB GWAS meta-analysis, we downloaded TGP EUR LD scores as a representative reference from LDSC. To match the SNP IDs to the rsIDs in the EUR LD scores, the GWAS summary statistics were annotated with dbSNP rsIDs v155 using the R package *SNPlocs.Hsapiens.dbSNP155.GRCh37*. The formatted summary statistics were used to compute observed-scale heritability for continuous traits and liability-scale heritability for binary traits. Sample prevalence for the four binary traits was estimated using phenotype data from the UK Biobank. Polygenicity for the ten traits was inferred using LDPred2-auto^51^. We used the precomputed EUR LD correlation matrix provided by LDPred2 and ran the Markov Chain Monte Carlo process with default hyperparameter settings.

### *All of Us* phenotype curation

For the 10 phenotypes with GWAS summary statistics available from Pan-UKBB, we curated the corresponding phenotypes in the *All of Us* following the methodology outlined in Tsuo *et al*^52^. We curated the quantitative traits based on concept IDs, and binary traits using PheCode map v1.2, which facilitate the mapping of ICD codes into their corresponding phecodes. For quantitative phenotypes, we harmonized their units by adopting the most frequently observed unit as the reference standard. We further removed those individuals with values exceeding 5 standard deviations from the mean. Subsequently, we performed inverse-rank normal transformation to ensure normalization.

### Target sample selection

Samples with both genotyped array and WGS data were selected. In total, 95,562 individuals had both types of data in *All of Us* v6. Samples were excluded if more than 10% of genotypes were missing. Related samples were removed based on a kinship score > 0.1. Samples predicted to be of EUR (European, composed of Finnish and non-Finnish European), AMR (Latino/admixed American), or AFR (African) ancestry were selected based on an auxiliary genetically predicted ancestry file provided in *All of Us*. Briefly, the ancestry labels were inferred using the prediction probabilities from a random forest classifier, which was trained on the 16-dimenional PCA space of reference HGDP and TGP samples and autosomal variants, which obtained from gnomAD^14^. A summary table of sample sizes for the 10 curated phenotypes is provided in Supplementary Table 2.

### Target variant QC

The QC criteria were identical for both array and WGS variants in *All of Us* v6. First, only autosomal variants were retained. Variants with a call rate lower than 90% were excluded. Indels were removed, and only common biallelic SNPs with a MAF > 0.01 were retained. QC of the 1,824,517 array variants resulted in 975,876 SNPs, while QC of the 702,574,937 WGS variants resulted in 8,996,707 SNPs (Supplementary Table 4). We did not impute array variants in *All of Us* due to the high computational cost associated with cloud-based analysis. *All of Us* genotype data is only released unimputed, such that this is expected to be standard practice for users of this particular dataset. *All of Us* array data is genotyped with the Global Diversity Array, which was designed to integrate highly optimized multiethnic variants with a genome-wide scaffold, incorporating WGS data from over 1,000 individuals of African and/or American ancestry, reflecting the genetic diversity of participants in the *All of Us* Research Program. As such, comparing polygenic prediction performance across ancestral groups using array and WGS data from *All of Us* is particularly relevant.

### C+T construction

For C+T, QC’d array and WGS variants were first restricted to those overlapping with QC’d GWAS summary statistics to maximize variant overlap. The counts of intersecting array or WGS variants with the discovery GWAS are presented in Supplementary Table 4. LD was estimated using 3,202 unrelated samples from five continental populations in the TGP Phase 3 as the reference panel; first- and second-degree relatives were excluded. Reference variants were filtered to retain common biallelic SNPs (MAF > 0.01). LD clumping was performed using PLINK to retain approximately independent variants (--clump-p1 1, --clump-r2 0.1, and --clump-kb 250). For each trait, clumped variants were subsequently filtered across a range of GWAS *p*-value thresholds (1, 0.5, 1e-1, 1e-2, 1e-3, 1e-4, 1e-5, 1e-6, 1e-7, and 1e-8), constituting the thresholding step of the C+T procedure. The number of retained variants after clumping is reported in Supplementary Table 5. PGS were calculated for each individual as:

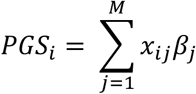

where *X* represents the dosage of alternate alleles {0, 1, 2} from the genotype matrix, *β* denotes the effect size estimates, *i* indexes the individual, *j* indexes the variant, and *M* is the total number of variants used for the calculation (i.e. number of clumped SNPs). Optimal *p*-value thresholds were selected separately for each population using 10-fold cross-validation (1 fold for validation and 9 folds for testing). Model performance was assessed using R^2^ for continuous traits and Nagelkerke’s R^2^ for binary traits. R^2^ is the proportion of variance explained by the full model relative to the null model, calculated using linear regression. Nagelkerke’s R^2^ was computed via logistic regression by comparing the likelihoods of the null and full models, adjusting for the overall fit to obtain a value indicating the proportion of variance explained by the PGS in the context of the full model. The null model included age, male sex, female sex, and the first 16 principal components as predictors, while the full model included all variables from the null model in addition to PGS.

### PRS-CS construction

We prepared the LD reference panel from two sources: (1) A precomputed EUR LD panel constructed using UKBB data, restricted to 1,120,696 HapMap3 variants, which was downloaded from the PRS-CS GitHub repository. (2) A genome-wide EUR LD panel constructed using the TGP and HGDP joint call set^53^. To generate this LD panel, we downloaded precomputed EUR LD blocks based on the GRCh38 genome from LDblocks_GRCh38^54^. We then computed the LD matrices for each LD block using PLINK, with a total of 7,330,169 post-QC (bi-allelic SNPs with MAF >0.5%) genome-wide variants from the VCF of 664 unrelated EUR samples in the TGP and HGDP joint call set. The LD matrices across 22 chromosomes were saved in HDF5 format, which is compatible with PRS-CS. To run PRS-CS, we did not set the global shrinkage parameter phi and allowed the program to automatically search for the optimal value. After obtaining posterior effect sizes generated from PRS-CS-auto, we computed PGS for each target individual by aggregating the product of their genotypes weighted by the updated effect sizes, as shown in the equation below:

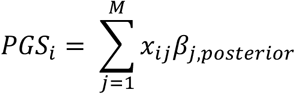

PGS performance was then evaluated using R^2^ for continuous traits and Nagelkerke’s R^2^ for binary traits, as described in the C+T section.

### Evaluation of prediction performances using PGS Catalog

To evaluate the PGS performance in the *AoU* and UKBB cohorts, we downloaded publicly available scoring files from the PGS Catalog for 4 traits: Height, HDL cholesterol, Type 2 Diabetes, and breast cancer. Each scoring file included SNP ID, effect allele, and corresponding effect size estimates derived from the published PGS models. The PGS studies selected were prioritized to those trained the model on multi-ancestry datasets and evaluated across multiple populations. Details of the selected scoring files are provided in Supplementary Table 13. Variant QC criteria for *AoU* data have been described previously. For UKBB array and imputed genotype, variants were restricted to bi-allelic SNPs with MAF > 0.005. Duplicate and ambiguous SNPs were removed, and alleles were harmonized between scoring files and the target genotypes, including strand and sign flipping where necessary. Phenotype curation for *AoU* was described previously. For UKBB, continuous traits were curated using the first data instance and values were converted with inverse rank normal transformation for each population groups (EUR, AMR, AFR, East Asian (EAS), and Central South Asian (CSA)). Binary disease traits were defined based on primary and secondary ICD-10 diagnosis codes, with individuals having at least one relevant code classified as cases and those without any diagnosis classified as controls. Polygenic scores were computed using PLINK to aggregate the weighted genotype dosage, where weights corresponded to effect size estimates from the PGS Catalog and genotypes were from *AoU*’s array and WGS data and from UKBB’s array and imputation data. Prediction performance was assessed with either R^2^ or Negelkerke R^2^, adjusting for age, sex, and 16 PCs.

### Simulation

We simulated phenotypes based on real genotype data from the *AoU* (array and WGS) and UKBB (array and imputation). For *AoU*, we randomly selected 50,000 unrelated EUR samples. To ensure comparable genotypes between array and WGS, we extracted QC’d chromosome 22 variants from the short-read WGS (srWGS) common Allele Count/Allele Frequency (ACAF) threshold callset (*AoU* release v7) to created two datasets: a full WGS variant set and an array-matched subset obtained by restricting WGS variants to those overlapping array SNPs. Phenotypes and effect sizes were simulated using GCTA^55^. Continuous phenotypes were simulated under 10 scenarios, each with 10 replicates, where each scenario differed in the proportion of causal variants present in array vs. WGS data (ranging from 100% to 0%) (Fig. 1b). Causal SNPs were randomly assigned to variants with allele frequencies between 0.005 and 0.05 across all scenarios. Two levels of genetic architecture were considered: 500 causal SNPs to represent polygenic traits (∼0.1% of the studied genomic variants) and 50 causal variants to represent sparse traits (∼0.01% of the studied genomic variants). Effect sizes were estimated by conducting GWAS on 80% of the samples with PGS computed on the remaining 20%. Ten PCs were generated for the discovery and target sets and included as covariates in GWAS and PGS analyses. For UKBB, we similarly selected 50,000 unrelated EUR samples and extracted chromosome 22 variants from the QC’d imputation data, subsequently generating an array-matched subset by restricting variants to those overlapping with *AoU* array SNPs to facilitate comparison. All other simulation and analysis procedures were identical to those applied to *AoU* data. PGS were constructed using C+T, since PRS-CS is limited to HapMap3 variants, which reduces the total number of SNPs. Predictive performance was evaluated using R^2^.

### Statistical fine-mapping

To identify likely causal variants in real data, we performed statistical fine-mapping using SuSiE^56^(susieR package v0.15.2), a Bayesian regression framework that models multiple causal variants within a locus. Pan-UKBB summary statistics (height, HDL, T2D, and breast cancer) and a precomputed UKBB EUR LD matrices were used as inputs. For each genome-wide significant locus (*p* < 5e-8), SuSiE was applied assuming up to 10 causal variants. Variants within 95% credible sets were prioritized, and those with posterior inclusion probabilities (PIPs) > 0.9 were considered likely causal for downstream analyses.

## Declaration of interests

A.R.M. has received speaker fees from Novartis. Other authors declare no competing interests.

## Acknowledgements

We gratefully acknowledge *All of Us* participants for their contributions, without whom this research would not have been possible. We also thank the National Institutes of Health’s *All of Us* Research Program for making available the participant genotype and phenotype data examined in this study. We thank Jessica Honorato Mauer for her assistance with QC of the UKBB imputed data and Aishi Ayyanathan for her help in generating the QC’d sample list used in this study. This research has been conducted using the UK Biobank Resource under Application Number 95179. This work was supported by NIH grants K01MH121659 and R01HG012869 to E.G.A.

## Author contributions

Y-S.L. analyzed the data, interpreted the results, and drafted the manuscript. T.T. analyzed the data and interpreted the results. Y.W. curated *All of Us* phenotypes. B.P. and A.R.M. provided critical comments. E.G.A designed and supervised the study and revised the manuscript. All authors approved the final manuscript.

## Web resources

Pan-UKBB Downloads page, https://pan.ukbb.broadinstitute.org/downloads

EUR LD scores, https://zenodo.org/records/8182036

TGP Phase 3 reference panels, https://www.cog-genomics.org/plink/2.0/resources

PRS-CS, https://github.com/getian107/PRScs

PGS Catalog, https://www.pgscatalog.org/

susieR, https://stephenslab.github.io/susieR/

UKBB LD reference, https://uchicago.app.box.com/s/jqocacd2fulskmhoqnasrknbt59x3xkn

## Data and code availability

The GWAS summary statistics used in this study were downloaded from the Pan UKBB Downloads page. Individual genotype data were obtained from the *All of Us* Research Program Controlled Tier Dataset v6. EUR LD scores from TGP, used for heritability estimation, were downloaded from the zenodo web resources describe above. The TGP Phase 3 reference panels used for C+T were downloaded from PLINK resources. The EUR reference LD panel, constructed using UKBB data for PRS-CS, was downloaded from the PRS-CS GitHub repository. PGS Catalog scoring files were downloaded from the link provided in Supplementary Table 13. The UKBB LD reference were downloaded from the UChicagoBox. All code used for this analysis is available at https://github.com/Atkinson-Lab/PGS_Array_WGS.

## Notes

https://github.com/Atkinson-Lab/PGS_Array_WGS

### Summary of Updates

The revised version includes comprehensive benchmarking of prediction models from the PGS Catalog, allowing for a more thorough comparison of their performance across genomic platforms. Additionally, it presents new results from both simulation studies and fine-mapping analyses, providing insights into the differential capture of causal variants and their impact on prediction accuracy across these platforms.

