## Supplementary Information for "Causal variant capture in genotype discovery approaches drives polygenic prediction performance across traits and populations"

### Supplementary Figures

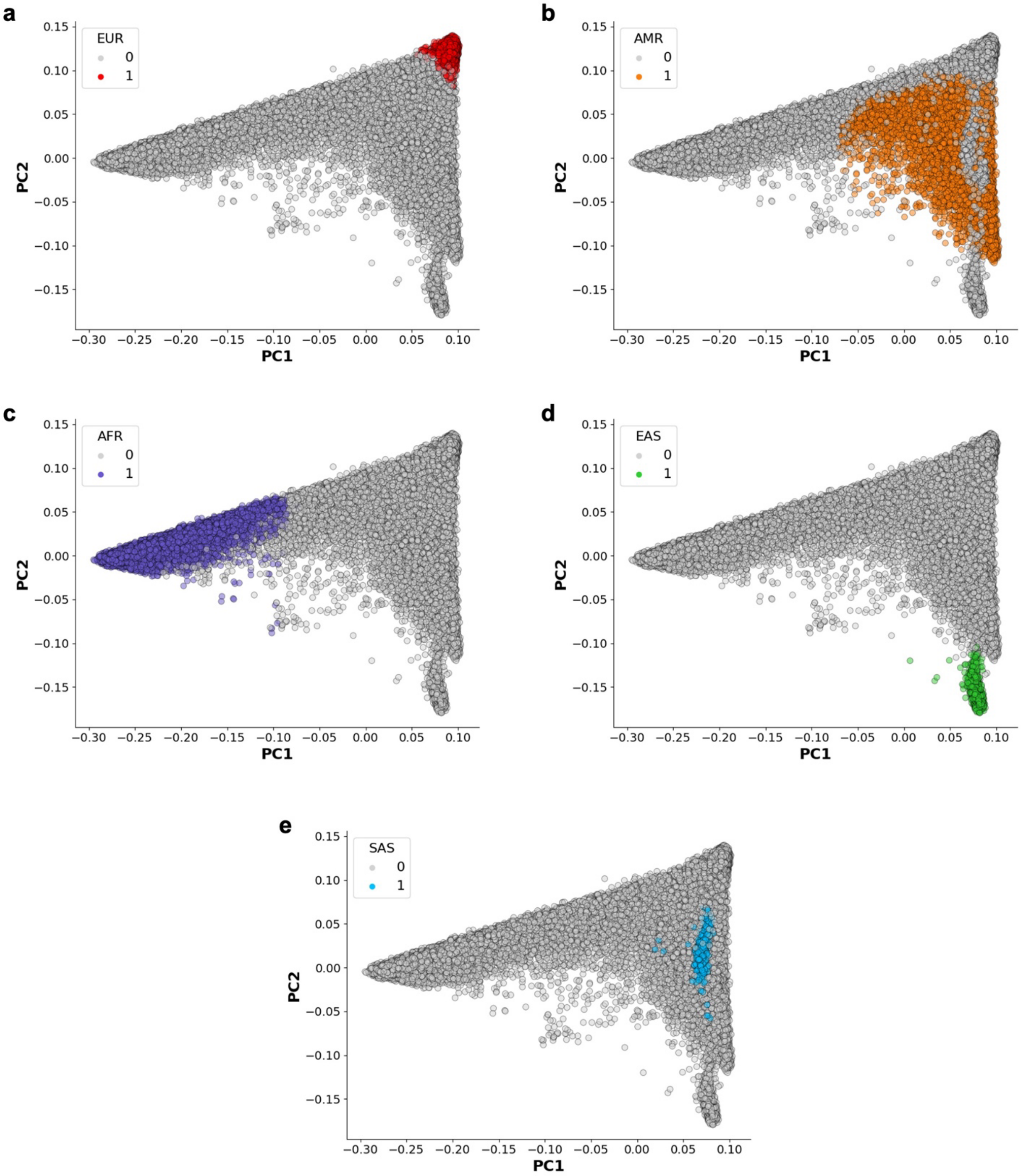


**Supplementary Fig. 1: PC plots for five populations in *All of Us*.**

The inferred genetic ancestries and PC values were downloaded from *All of Us*. Individuals clustered by PC values are shown for (a) EUR, (b) AMR, (c) AFR, (d) EAS, and (e) SAS.

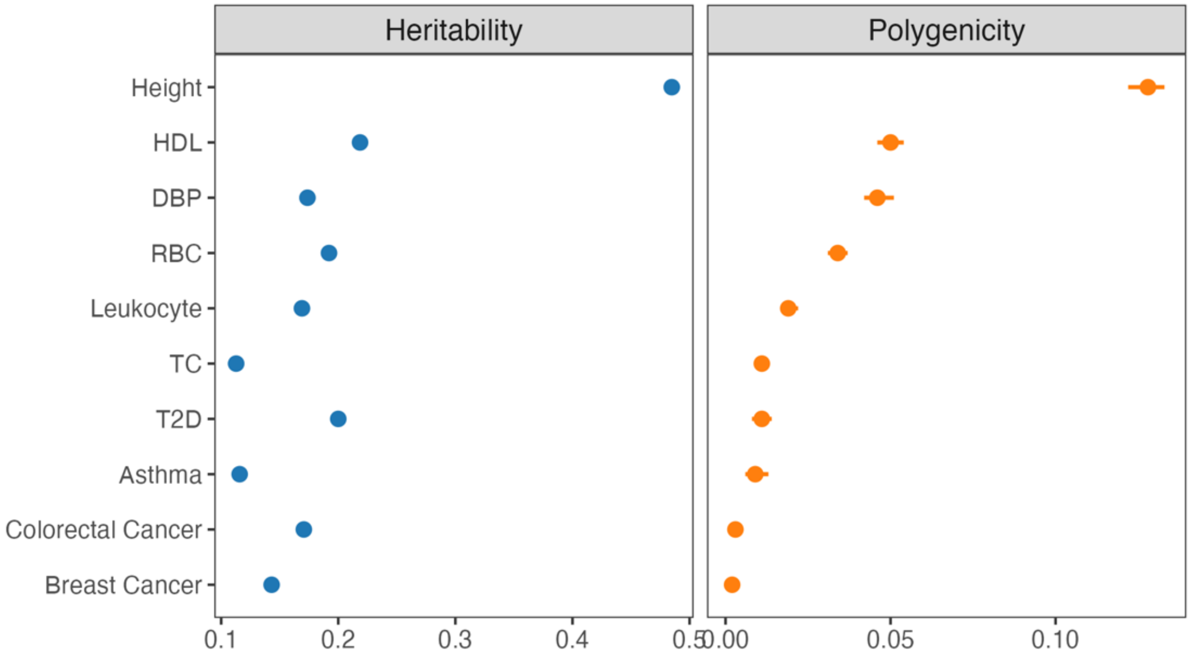


**Supplementary Fig. 2: SNP heritability and polygenicity for ten traits in EUR.**

Left: Observed-scale heritability for the six continuous traits and liability-scale heritability for the four binary traits from Pan-UKBB, estimated using LDSC. Right: Polygenicity inferred using LDPred2-auto. Traits are ordered by polygenicity.

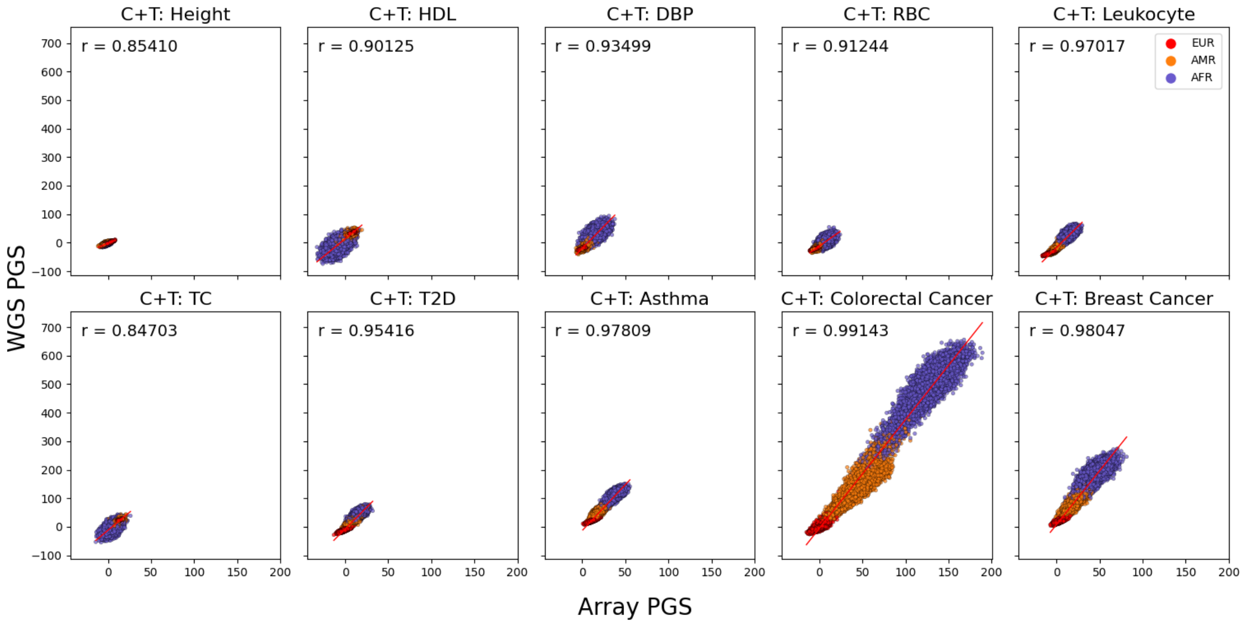


**Supplementary Fig. 3**: **Correlation between Array PGS sand WGS PGS using C+T.**

For each sample, the average scores from 10-fold cross-validation calculated using C+T are plotted. Samples are colored by population. The correlation coefficient (r) between the Array PGS and WGS PGS is shown. Note the larger spread for admixed populations.

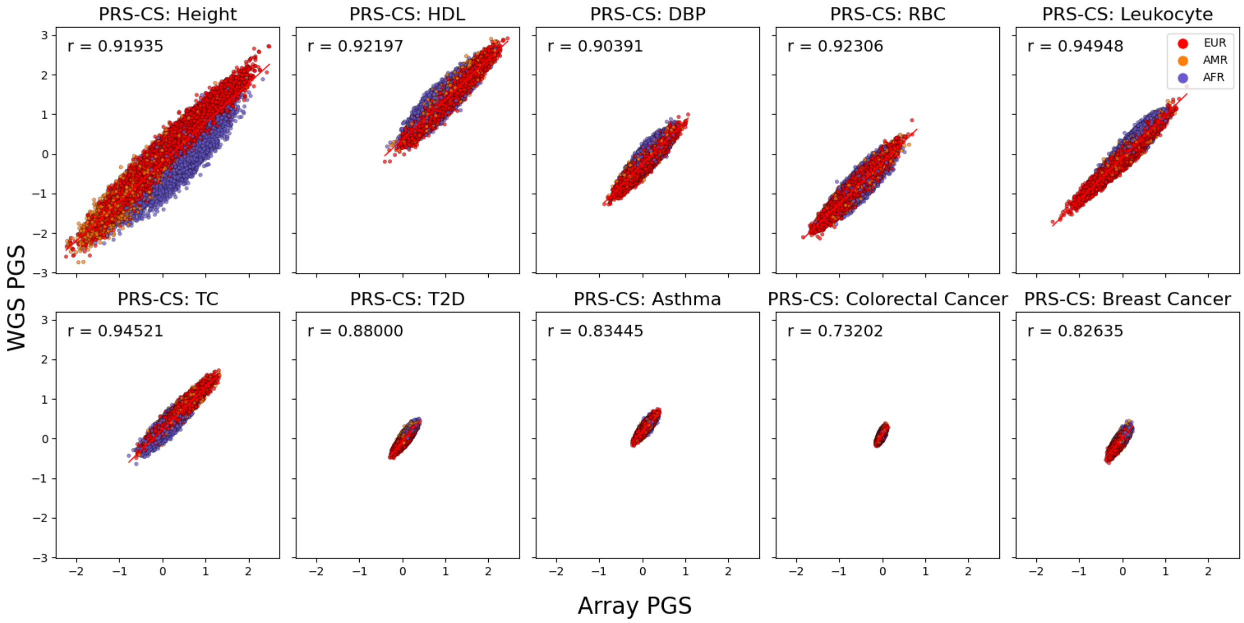


**Supplementary Fig. 4: Correlation between Array PGS and WGS PGS using PRS-CS.**

For each sample, the score calculated using PRS-CS was plotted. Samples are colored by population. The correlation coefficient (r) between the Array PGS and WGS PGS is shown.

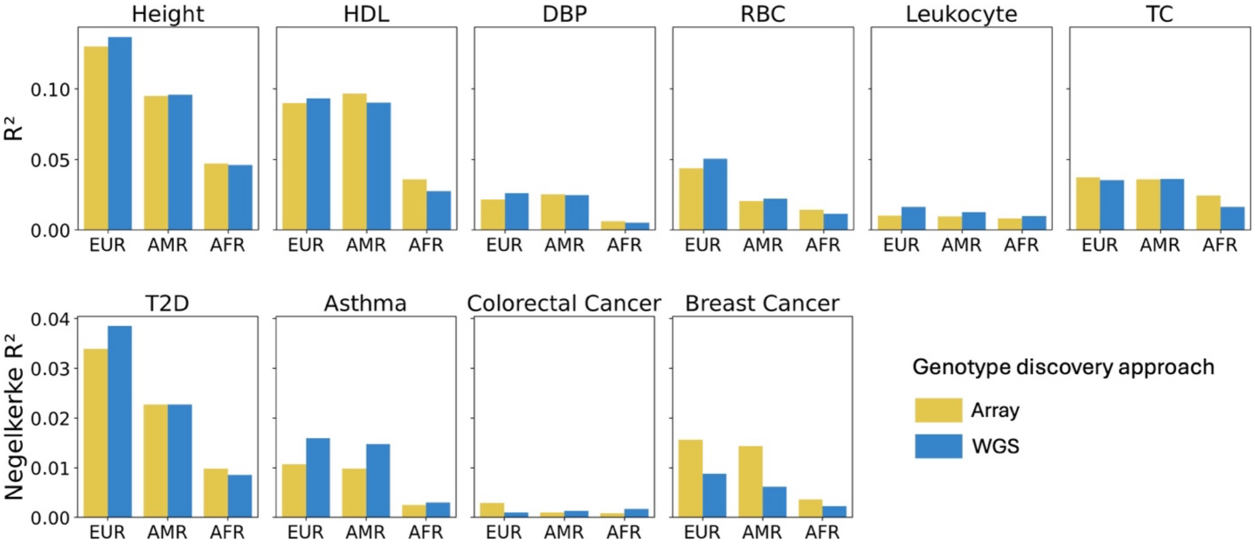


**Supplementary Fig. 5: Comparison of PGS performance across genotype discovery approaches using PRS-CS with full-scale LD panel.**

PGS were computed for samples with array- and WGS-genotypes in *All of Us*. $R^2$ or Nagelkerke's $R^2$ were determined by adjusting for age, sex, and the first 16 genetic PCs.

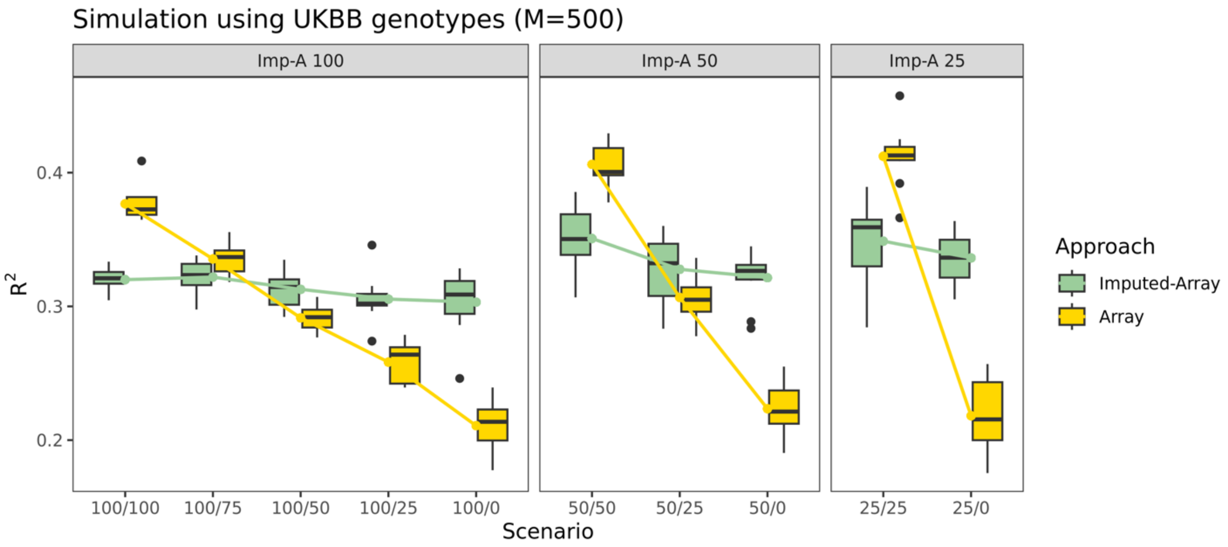


**Supplementary Fig. 6: Simulation settings of causal variants captured in UKBB.**

Traits were simulated using UKBB imputation genotypes under polygenic settings ($M$ = 500 causal variants) with heritability set to 0.5. Let $P_{imputation}$ denote the percentage of causal variants captured by the imputed-array, and $P_{array}$ denote the percentage captured by the array. Each simulation scenario is labeled as $P_{imputation}$ / $P_{array}$. Imp-A: Imputed-Array.

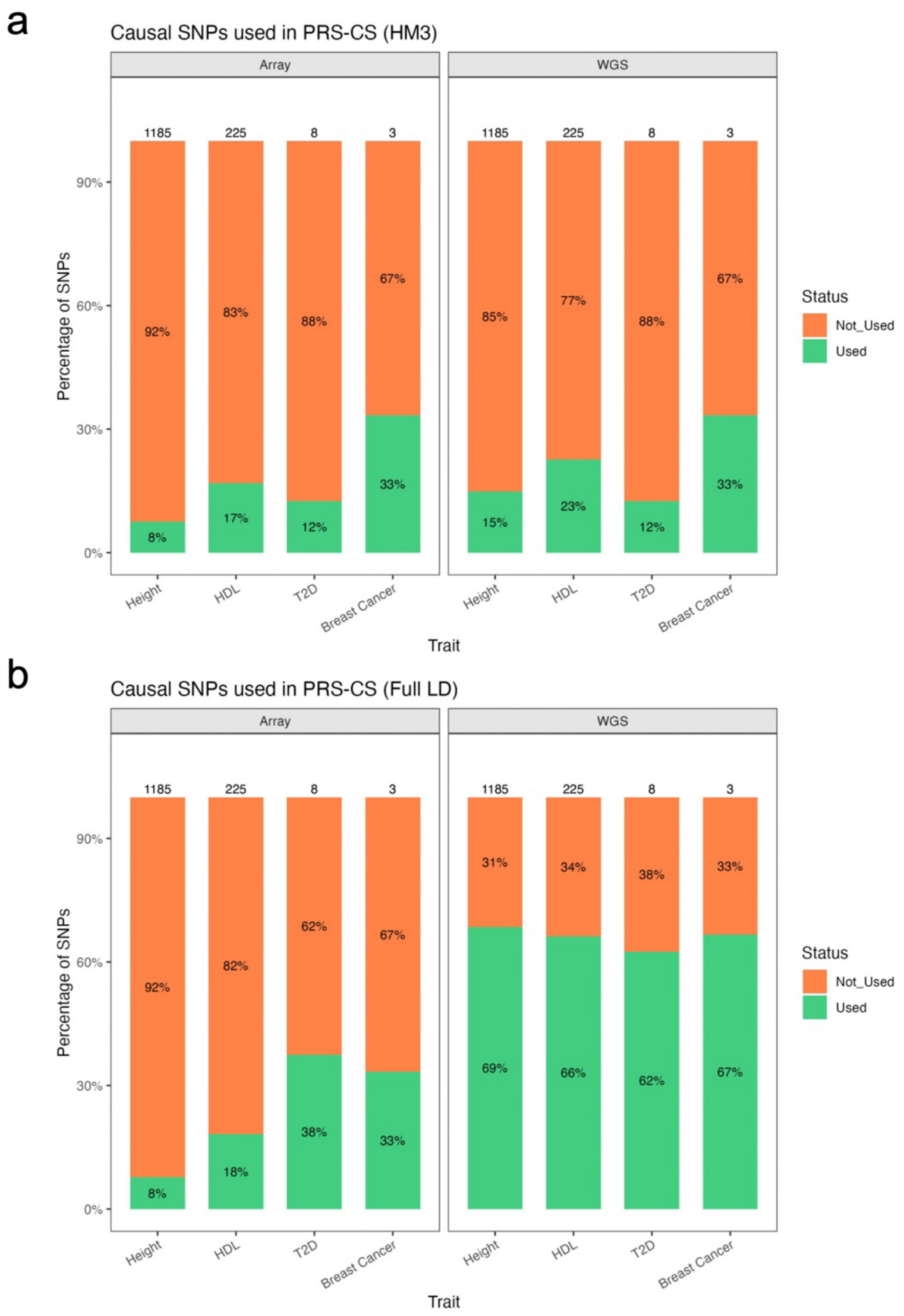


**Supplementary Fig. 7: Proportion of likely causal SNPs captured by genotype discovery approaches.**

Bar graphs show the percentage of likely causal SNPs in PRS-CS, overlapped with (a) the HapMap3 LD panel and (b) the full-scale LD panel. The total number of inferred causal SNPs is labeled above each bar. The orange portion represents the percentage of causal SNPs not captured by the genotype discovery approach, while the green portion represents the percentage that were captured.

### Supplementary Tables

#### Supplementary Table 1. Sample size for Pan-UKBB traits by population.

| | Height | HDL | DBP | RBC | Leukocyte | TC | T2D | Asthma | Colorectal cancer | Breast cancer |
|---|---|---|---|---|---|---|---|---|---|---|
| **Phecode** | 50 | 30760 | DBP | 30010 | 30000 | 30690 | 250.2 | 495 | 153 | 250.2 |
| **Description** | Standing height | HDL cholesterol | Diastolic blood pressure, automated reading, adjusted by medication | Red blood cell (erythrocyte ) count | White blood cell (leukocyte) count | Cholesterol | Type 2 diabetes | Asthma | Colorectal cancer | Type 2 diabetes |
| **Cases, N** | 438,478 | 385,023 | 413,288 | 427,938 | 427,933 | 420,607 | 25,550 | 33,000 | 5,807 | 13,631 |
| **Cases hq, N** | 434,809 | 374,709 | 402,068 | 418,075 | 416,522 | 409,385 | 24,432 | - | - | - |
| **AFR** | 6,556 | 5,754 | 6,350 | 6,281 | 6,281 | 6,212 | 796 | 576 | 64 | 133 |
| **AMR** | 972 | 854 | 930 | 950 | 950 | 938 | - | 64 | - | - |
| **CSA** | 8,657 | 7,688 | 8,206 | 8,532 | 8,532 | 8,422 | 1,664 | 903 | 50 | 163 |
| **EAS** | 2,697 | 2,342 | 2,466 | 2,632 | 2,632 | 2,572 | 152 | 149 | - | 78 |
| **EUR** | 419,596 | 367,021 | 393,862 | 407,995 | 407,990 | 400,963 | 22,768 | 311,69 | 5,693 | 13,257 |
| **MID** | - | 1,364 | 1,474 | 1,548 | 1,548 | 1,500 | 170 | 139 | - | - |
| **Controls, N** | - | - | - | - | - | - | 413,134 | 398,413 | 401,350 | 214,950 |
| **AFR** | | | | | | | 5,793 | 5,999 | 6,260 | 3,613 |
| **AMR** | | | | | | | - | 898 | - | - |
| **CSA** | | | | | | | 7,181 | 7,886 | 8,350 | 3,774 |
| **EAS** | | | | | | | 2,555 | 2,540 | - | 1,650 |
| **EUR** | | | | | | | 396,181 | 379,656 | 386,740 | 205,913 |
| **MID** | | | | | | | 1,424 | 1,434 | - | - |

Table showing total sample size and sample size for each population. AFR: African; AMR: Admixed American; CSA: Central/South Asian; EAS: East Asian; EUR: European; MID: Middle Eastern.

**Supplementary Table 2. Sample size for *All of Us* traits by population.**

| | Height | HDL | DBP | RBC | Leukocyte | TC | T2D | Asthma | Colorectal cancer | Breast cancer |
|---|---|---|---|---|---|---|---|---|---|---|
| **Cases, N** | 90,521 | 36,721 | 91,231 | 50,003 | 48,399 | 37,720 | 8,041 | 11,384 | 766 | 2,674 |
| **EUR** | 45,348 | 21,575 | 45,741 | 27,410 | 26,345 | 22,165 | 3,714 | 6,137 | 502 | 1,761 |
| **AMR** | 13,448 | 4,264 | 13,612 | 7,087 | 6,513 | 4,430 | 1,360 | 1,533 | 74 | 245 |
| **AFR** | 20,527 | 6,330 | 20,599 | 9,147 | 9,349 | 6,483 | 2,114 | 2,306 | 90 | 320 |
| **EAS** | 1,996 | 669 | 2,002 | 863 | 801 | 676 | 97 | 139 | 13 | 51 |
| **SAS** | 923 | 373 | 929 | 432 | 418 | 377 | 87 | 54 | 2 | 10 |
| **Controls, N** | - | - | - | - | - | - | 82,232 | 51,461 | 90,864 | 51,097 |
| **EUR** | | | | | | | 41,355 | 22,794 | 45,218 | 24,903 |
| **AMR** | | | | | | | 12,146 | 8,687 | 13,668 | 8,732 |
| **AFR** | | | | | | | 18,349 | 13,331 | 20,689 | 11,104 |
| **EAS** | | | | | | | 1,910 | 1,466 | 1,232 | 2,009 |
| **SAS** | | | | | | | 838 | 649 | 923 | 443 |

Table showing total sample size (N) and sample size for each population.

**Supplementary Table 3. Numerical values of heritability and polygenicity for EUR.**

| ***Trait*** | **Heritability** | **Polygenicity [95%CI]** |
|---|---|---|
| *Height* | 0.4848 | 0.128 [0.122, 0.133] |
| *HDL* | 0.2186 | 0.05 [0.046, 0.054] |
| *DBP* | 0.1737 | 0.046 [0.042, 0.051] |
| *RBC* | 0.1921 | 0.034 [0.031, 0.037] |
| *Leukocyte* | 0.169 | 0.019 [0.017, 0.022] |
| *TC* | 0.1128 | 0.011 [0.009, 0.012] |
| *T2D* | 0.2 | 0.011 [0.008, 0.014] |
| *Asthma* | 0.1158 | 0.009 [0.006, 0.013] |
| *Colorectal Cancer* | 0.1706 | 0.003 [0.001, 0.005] |
| *Breast Cancer* | 0.1431 | 0.002 [0.001, 0.003] |

Observed-scale heritability for the six continuous traits and liability-scale heritability for the four binary traits from Pan-UKBB, estimated using LDSC. Polygenicity inferred using LDPred2-auto. Traits are ordered by polygenicity.

**Supplementary Table 4. Overlap between array or WGS variants and variants identified in the discovery GWAS.**

| ***Trait*** | | ***Array*** | ***WGS*** | **GWAS** |
|---|---|---|---|---|
| | (Pre-QC) | 1,824,517 | 702,574,937 | |
| | (Post-QC) | 975,876 | 8,996,707 | |
| | **GWAS ∩ target** | | | |
| *Height* | | 918,553 | 8,604,715 | 28,987,534 |
| *HDL* | | 914,816 | 8,484,538 | 28,987,534 |
| *DBP* | | 915,419 | 8,503,485 | 28,987,534 |
| *RBC* | | 916,315 | 8,533,326 | 28,987,534 |
| *Leukocyte* | | 915,594 | 8,509,546 | 28,987,534 |
| *TC* | | 915,512 | 8,506,131 | 28,987,534 |
| *T2D* | | 918,526 | 8,604,426 | 28,987,534 |
| *Asthma* | | 918,610 | 8,605,299 | 28,987,534 |
| *Colorectal cancer* | | 915,045 | 8,537,841 | 28,987,534 |
| *Breast cancer* | | 918,242 | 8,601,970 | 28,987,534 |

Values represent variant counts.

**Supplementary Table 5. Array and WGS variant count after LD clumping by trait.**

| ***Trait*** | **Clumped variants (% reduction)** | |
|---|---|---|
| | *Array* | *WGS* |
| *Height* | 262,399 (71.43) | 468,717 (94.55) |
| *HDL* | 261,583 (71.41) | 465,726 (94.51) |
| *DBP* | 261,858 (71.39) | 466,453 (94.51) |
| *RBC* | 262,290 (71.38) | 466,830 (94.53) |
| *Leukocyte* | 261,711 (71.42) | 466,426 (94.52) |
| *TC* | 262,106 (71.37) | 466,573 (94.51) |
| *T2D* | 262,905 (71.38) | 466,573 (94.58) |
| *Asthma* | 262,804 (71.39) | 468,426 (94.56) |
| *Colorectal cancer* | 261,535 (71.42) | 464,134 (94.56) |
| *Breast cancer* | 262,969 (71.36) | 468,563 (94.55) |

Parentheses indicate the proportion of variants removed after clumping, relative to the intersection of target and GWAS variants from the previous step.

**Supplementary Table 8. Comparison of polygenic prediction across traits and genotype discovery approaches using C+T and PRS-CS.**

| ***Trait*** | **Population** | **$R^2$** | | | |
|---|---|---|---|---|---|
| | | ***C+T*** | | ***PRS-CS*** | |
| | | **Array (95% CI)** | **WGS (95% CI)** | **Array** | **WGS** |
| *Height* | EUR | $9.95\times10^{-2}$ ($9.92\times10^{-2}$, $9.98\times10^{-2}$) | $1.07\times10^{-1}$ ($1.07\times10^{-1}$, $1.08\times10^{-1}$) | $1.34\times10^{-1}$ | $1.45\times10^{-1}$ |
| | AMR | $5.22\times10^{-2}$ ($5.14\times10^{-2}$, $5.31\times10^{-2}$) | $5.34\times10^{-2}$ ($5.15\times10^{-2}$, $5.53\times10^{-2}$) | $9.45\times10^{-2}$ | $1.04\times10^{-1}$ |
| | AFR | $3.14\times10^{-2}$ ($3.02\times10^{-2}$, $3.27\times10^{-2}$) | $3.37\times10^{-2}$ ($3.30\times10^{-2}$, $3.45\times10^{-2}$) | $4.82\times10^{-2}$ | $5.68\times10^{-2}$ |
| *HDL* | EUR | $5.44\times10^{-2}$ ($5.35\times10^{-2}$, $6.68\times10^{-2}$) | $5.28\times10^{-2}$ ($5.07\times10^{-2}$, $5.49\times10^{-2}$) | $9.31\times10^{-2}$ | $1.02\times10^{-1}$ |
| | AMR | $6.14\times10^{-2}$ ($5.59\times10^{-2}$, $5.59\times10^{-2}$) | $4.81\times10^{-2}$ ($4.55\times10^{-2}$, $5.06\times10^{-2}$) | $9.83\times10^{-2}$ | $1.02\times10^{-1}$ |
| | AFR | $2.73\times10^{-2}$ ($2.60\times10^{-2}$, $2.86\times10^{-2}$) | $8.86\times10^{-3}$ ($5.68\times10^{-3}$, $1.20\times10^{-2}$) | $3.27\times10^{-2}$ | $3.81\times10^{-2}$ |
| *DBP* | EUR | $1.12\times10^{-2}$ ($1.02\times10^{-2}$, $1.22\times10^{-2}$) | $1.02\times10^{-2}$ ($9.47\times10^{-3}$, $1.10\times10^{-2}$) | $2.16\times10^{-2}$ | $2.54\times10^{-2}$ |
| | AMR | $5.43\times10^{-3}$ ($4.50\times10^{-3}$, $6.37\times10^{-3}$) | $4.78\times10^{-3}$ ($3.94\times10^{-3}$, $5.61\times10^{-3}$) | $2.37\times10^{-2}$ | $2.68\times10^{-2}$ |
| | AFR | $1.36\times10^{-3}$ ($9.66\times10^{-4}$, $1.75\times10^{-3}$) | $1.37\times10^{-3}$ ($8.80\times10^{-4}$, $1.87\times10^{-3}$) | $5.53\times10^{-3}$ | $8.13\times10^{-3}$ |
| *RBC* | EUR | $2.72\times10^{-2}$ ($2.63\times10^{-2}$, $2.80\times10^{-2}$) | $2.81\times10^{-2}$ ($2.73\times10^{-2}$, $2.89\times10^{-2}$) | $4.75\times10^{-2}$ | $5.20\times10^{-2}$ |
| | AMR | $9.24\times10^{-3}$ ($7.33\times10^{-3}$, $1.12\times10^{-2}$) | $1.10\times10^{-2}$ ($8.76\times10^{-3}$, $1.32\times10^{-2}$) | $2.11\times10^{-2}$ | $2.65\times10^{-2}$ |
| | AFR | $1.09\times10^{-2}$ ($8.85\times10^{-3}$, $1.29\times10^{-2}$) | $6.82\times10^{-3}$ ($4.28\times10^{-3}$, $9.35\times10^{-3}$) | $1.43\times10^{-2}$ | $1.55\times10^{-2}$ |
| *Leukocyte* | EUR | $9.83\times10^{-3}$ ($9.53\times10^{-3}$, $1.01\times10^{-2}$) | $1.06\times10^{-2}$ ($1.02\times10^{-2}$, $1.10\times10^{-2}$) | $1.55\times10^{-2}$ | $1.90\times10^{-2}$ |
| | AMR | $7.73\times10^{-3}$ ($5.86\times10^{-3}$, $9.60\times10^{-3}$) | $6.61\times10^{-3}$ ($6.17\times10^{-3}$, $7.05\times10^{-3}$) | $1.22\times10^{-2}$ | $1.59\times10^{-2}$ |
| | AFR | $8.89\times10^{-3}$ ($7.54\times10^{-3}$, $1.02\times10^{-2}$) | $1.02\times10^{-2}$ ($9.76\times10^{-3}$, $1.06\times10^{-2}$) | $1.36\times10^{-2}$ | $1.45\times10^{-2}$ |
| *TC* | EUR | $2.14\times10^{-2}$ ($2.10\times10^{-2}$, $2.18\times10^{-2}$) | $2.35\times10^{-2}$ ($2.29\times10^{-2}$, $2.42\times10^{-2}$) | $4.08\times10^{-2}$ | $4.27\times10^{-2}$ |
| | AMR | $2.02\times10^{-2}$ ($1.74\times10^{-2}$, $2.31\times10^{-2}$) | $2.09\times10^{-2}$ ($1.99\times10^{-2}$, $2.19\times10^{-2}$) | $4.46\times10^{-2}$ | $4.33\times10^{-2}$ |
| | AFR | $7.55\times10^{-3}$ ($5.02\times10^{-3}$, $1.01\times10^{-2}$) | $5.02\times10^{-3}$ ($2.10\times10^{-3}$, $7.95\times10^{-3}$) | $2.34\times10^{-2}$ | $2.20\times10^{-2}$ |
| *T2D* | EUR | $2.29\times10^{-2}$ ($2.17\times10^{-2}$, $2.40\times10^{-2}$) | $2.05\times10^{-2}$ ($2.01\times10^{-2}$, $2.10\times10^{-2}$) | $3.65\times10^{-2}$ | $4.41\times10^{-2}$ |
| | AMR | $8.50\times10^{-3}$ ($7.04\times10^{-3}$, $9.96\times10^{-3}$) | $4.88\times10^{-3}$ ($4.44\times10^{-3}$, $5.23\times10^{-3}$) | $2.21\times10^{-2}$ | $2.90\times10^{-2}$ |
| | AFR | $3.83\times10^{-3}$ ($2.99\times10^{-3}$, $5.67\times10^{-3}$) | $3.74\times10^{-3}$ ($3.30\times10^{-3}$, $4.18\times10^{-3}$) | $1.08\times10^{-2}$ | $1.09\times10^{-2}$ |
| *Asthma* | EUR | $1.08\times10^{-2}$ ($8.97\times10^{-3}$, $1.25\times10^{-2}$) | $1.04\times10^{-2}$ ($9.96\times10^{-3}$, $1.08\times10^{-2}$) | $1.39\times10^{-2}$ | $1.77\times10^{-2}$ |
| | AMR | $4.71\times10^{-3}$ ($3.70\times10^{-3}$, $5.72\times10^{-3}$) | $2.55\times10^{-3}$ ($2.11\times10^{-3}$, $2.99\times10^{-3}$) | $1.15\times10^{-2}$ | $1.74\times10^{-2}$ |
| | AFR | $6.82\times10^{-4}$ ($3.70\times10^{-4}$, $9.95\times10^{-4}$) | $1.39\times10^{-3}$ ($9.53\times10^{-4}$, $1.83\times10^{-3}$) | $3.33\times10^{-3}$ | $3.67\times10^{-3}$ |
| *Colorectal Cancer* | EUR | $2.41\times10^{-3}$ ($1.53\times10^{-3}$, $3.28\times10^{-3}$) | $1.58\times10^{-3}$ ($1.14\times10^{-3}$, $2.02\times10^{-3}$) | $2.60\times10^{-3}$ | $1.37\times10^{-3}$ |
| | AMR | $1.36\times10^{-3}$ ($-8.07\times10^{-5}$, $2.80\times10^{-3}$) | $3.81\times10^{-4}$ ($-5.61\times10^{-5}$, $8.19\times10^{-4}$) | $3.10\times10^{-3}$ | $1.50\times10^{-3}$ |
| | AFR | $1.44\times10^{-3}$ ($-3.52\times10^{-5}$, $2.91\times10^{-3}$) | $1.08\times10^{-3}$ ($6.44\times10^{-4}$, $1.52\times10^{-3}$) | $1.67\times10^{-4}$ | $4.80\times10^{-4}$ |
| *Breast Cancer* | EUR | $1.17\times10^{-2}$ ($1.06\times10^{-2}$, $1.28\times10^{-2}$) | $8.02\times10^{-3}$ ($7.59\times10^{-3}$, $8.46\times10^{-3}$) | $1.57\times10^{-2}$ | $1.30\times10^{-2}$ |
| | AMR | $2.93\times10^{-3}$ ($1.56\times10^{-3}$, $4.31\times10^{-3}$) | $2.77\times10^{-3}$ ($2.33\times10^{-3}$, $3.21\times10^{-3}$) | $1.41\times10^{-2}$ | $1.31\times10^{-2}$ |
| | AFR | $1.07\times10^{-3}$ ($3.31\times10^{-4}$, $1.81\times10^{-3}$) | $7.33\times10^{-4}$ ($2.96\times10^{-4}$, $1.17\times10^{-3}$) | $4.11\times10^{-3}$ | $3.93\times10^{-3}$ |

The $R^2$ columns report both $R^2$ and pseudo-$R^2$. For C+T, the values represent the average of 10 $R^2$ values. Confidence intervals are shown in parentheses.

**Supplementary Table 9. Paired t-test results comparing the $R^2$ of array- and WGS-derived PGS using C+T.**

| *Trait* | Population | T-statistic | *P*-value | Significance |
|---|---|---|---|---|
| *Height* | EUR | -39.987126 | 1.90E-11 | *** |
| | AMR | -1.2313768 | 0.24938743 | ns |
| | AFR | -3.7645248 | 0.00445385 | ** |
| *HDL* | EUR | 1.68330555 | 0.12660573 | ns |
| | AMR | 5.57916618 | 0.00034332 | *** |
| | AFR | 11.9992443 | 7.70E-07 | *** |
| *DBP* | EUR | 4.46539144 | 0.00156503 | ** |
| | AMR | 1.52683755 | 0.16114689 | ns |
| | AFR | -0.0489786 | 0.96200609 | ns |
| *RBC* | EUR | -3.361988 | 0.00836271 | ** |
| | AMR | -1.2705147 | 0.23576005 | ns |
| | AFR | 3.39305722 | 0.00796043 | ** |
| *Leukocyte* | EUR | -8.480171 | 1.39E-05 | *** |
| | AMR | 2.41028038 | 0.03923098 | * |
| | AFR | -2.3803669 | 0.04120291 | * |
| *TC* | EUR | -11.101762 | 1.49E-06 | *** |
| | AMR | -0.5540333 | 0.59305281 | ns |
| | AFR | 1.47267495 | 0.17492715 | ns |
| *T2D* | EUR | 3.01761282 | 0.01453542 | * |
| | AMR | 7.75629299 | 2.83E-05 | *** |
| | AFR | 0.18786969 | 0.85514629 | ns |
| *Asthma* | EUR | 0.44537792 | 0.66656658 | ns |
| | AMR | 3.02458138 | 0.01437226 | * |
| | AFR | -2.8937323 | 0.01777588 | * |
| *Colorectal Cancer* | EUR | 1.94575126 | 0.08353466 | ns |
| | AMR | 1.45908043 | 0.17854408 | ns |
| | AFR | 0.4105244 | 0.69102257 | ns |
| *Breast Cancer* | EUR | 7.49041365 | 3.73E-05 | *** |
| | AMR | 0.14845857 | 0.88525396 | ns |
| | AFR | 1.05547058 | 0.31871845 | ns |

Paired t-tests were performed for each trait and each population. Fields highlighted in blue indicate that PGS using WGS significantly outperforms array genotype. Fields highlighted in yellow indicate that PGS using array significantly outperforms WGS genotype. Asterisks denote the level of significance: *$P < 0.05$; **$P < 0.01$; ***$P < 0.001$.

**Supplementary Table 10. Previous studies on selected traits in underrepresented populations.**

| *Trait* | Description | Associated variants | References | Effect size in Pan-UKBB $(-\log_{10}(p))$ |
|---|---|---|---|---|
| *Leukocyte* | Individuals of African descent have been shown to have lower white blood cell and neutrophil counts than European Americans | Duffy Null polymorphism rs2814778 | [1,2] | 18.06 |
| *Asthma* | Asthma is twice as prevalent in African American compared to European American individuals, likely due both to social determinants of health as well as unique genetic variation | Not indicated in the study | [3] | |
| *HDL* | Black/African American individuals tend to have higher levels of HDL cholesterol than white individuals | | [4,5] | |
| | A meta-analysis of 15,901 African Americans for HDL | rs868213 in EXOC3L1 and HDL | [6] | 10.29 |
| | Low HDL cholesterol was more prevalent in Hispanic adults than in non-Hispanic black and non-Hispanic white | | [7] | |
| *RBC* | GWAS of RBC count identified in African Americans | rs13335629 | [8] | 8.39 |
| *T2D* | The prevalence of T2D is higher in Hispanics/Latinos than in non-Hispanic whites | rs7903146 in TCF7L2 | [9,10] | 161.1 |
| | | rs2283228 in KCNQ1 | | 5.177 |

Note that the self-identified race and ethnicity list in the table may not be identical to the genetic ancestry (AFR and AMR) used in this study.

**Supplementary Table 11. Variants overlapping between the target dataset, GWAS, and the LD panel used in PRS-CS.**

| ***Trait*** | **HM3 ∩ GWAS ∩ target (% overlap)** | | **FS ∩ GWAS ∩ target (% overlap)** | |
|---|---|---|---|---|
| | *Array* | *WGS* | *Array* | *WGS* |
| *Height* | 284,596 (29.16) | 924,300 (10.27) | 685,764 (70.27) | 5,467,939 (60.78) |
| *HDL* | 284,596 (29.16) | 924,300 (10.27) | 685,759 (70.27) | 5,467,919 (60.78) |
| *DBP* | 284,596 (29.16) | 924,300 (10.27) | 685,761 (70.27) | 5,467,919 (60.78) |
| *RBC* | 284,596 (29.16) | 924,300 (10.27) | 685,762 (70.27) | 5,467,920 (60.78) |
| *Leukocyte* | 284,596 (29.16) | 924,300 (10.27) | 685,761 (70.27) | 5,467,918 (60.78) |
| *TC* | 284,596 (29.16) | 924,300 (10.27) | 685,761 (70.27) | 5,467,920 (60.78) |
| *T2D* | 284,596 (29.16) | 924,300 (10.27) | 685,775 (70.27) | 5,467,938 (60.78) |
| *Asthma* | 284,596 (29.16) | 924,300 (10.27) | 685,770 (70.27) | 5,436,862 (60.43) |
| *Colorectal cancer* | 284,594 (29.16) | 924,294 (10.27) | 685,760 (70.27) | 5,467,492 (60.77) |
| *Breast cancer* | 284,596 (29.16) | 924,300 (10.27) | 685,775 (70.27) | 5,467,916 (60.78) |

In the *All of Us* dataset, the intersection of the GWAS, QC'd array data, and the HapMap3 LD matrix (HM3 ∩ GWAS ∩ target) included approximately 0.3 million variants (~29% of the QC'd array variants), while the intersection with the full-scale LD matrix (FS ∩ GWAS ∩ target) comprised around 0.7 million variants (~70% of the QC'd array variants). For the QC'd WGS data, the intersection with the GWAS and the HapMap3 LD matrix (HM3 ∩ GWAS ∩ target) included approximately 0.9 million variants (~10% of the QC'd WGS variants), whereas the intersection with the full-scale LD matrix (FS ∩ GWAS ∩ target) contained about 5.5 million variants (~61% of the QC'd WGS variants).

Parentheses indicate the proportion of variants shared across all three sets (LD panel ∩ GWAS ∩ target), relative to the total in the target set. HM3: HapMap3 LD panel. FS: Full-scale LD panel.

**Supplementary Table 12. Comparison of polygenic prediction across traits and genotype discovery approaches using PRS-CS with a full-scale LD panel.**

| ***Trait*** | **Population** | **$R^2$** | |
|---|---|---|---|
| | | *Array* | *WGS* |
| *Height* | EUR | $1.30 \times 10^{-1}$ | $1.37 \times 10^{-1}$ |
| | AMR | $9.50 \times 10^{-2}$ | $9.59 \times 10^{-2}$ |
| | AFR | $4.70 \times 10^{-2}$ | $4.60 \times 10^{-2}$ |
| *HDL* | EUR | $8.99 \times 10^{-2}$ | $9.33 \times 10^{-2}$ |
| | AMR | $9.66 \times 10^{-2}$ | $9.03 \times 10^{-2}$ |
| | AFR | $3.59 \times 10^{-2}$ | $2.74 \times 10^{-2}$ |
| *DBP* | EUR | $2.15 \times 10^{-2}$ | $2.62 \times 10^{-2}$ |
| | AMR | $2.53 \times 10^{-2}$ | $2.48 \times 10^{-2}$ |
| | AFR | $6.29 \times 10^{-3}$ | $4.93 \times 10^{-3}$ |
| *RBC* | EUR | $4.36 \times 10^{-2}$ | $5.04 \times 10^{-2}$ |
| | AMR | $2.04 \times 10^{-2}$ | $2.23 \times 10^{-2}$ |
| | AFR | $1.44 \times 10^{-2}$ | $1.14 \times 10^{-2}$ |
| *Leukocyte* | EUR | $9.98 \times 10^{-3}$ | $1.62 \times 10^{-2}$ |
| | AMR | $9.68 \times 10^{-3}$ | $1.26 \times 10^{-2}$ |
| | AFR | $8.04 \times 10^{-3}$ | $9.91 \times 10^{-3}$ |
| *TC* | EUR | $3.73 \times 10^{-2}$ | $3.53 \times 10^{-2}$ |
| | AMR | $3.60 \times 10^{-2}$ | $3.62 \times 10^{-2}$ |
| | AFR | $2.44 \times 10^{-2}$ | $1.63 \times 10^{-2}$ |
| *T2D* | EUR | $3.39 \times 10^{-2}$ | $3.85 \times 10^{-2}$ |
| | AMR | $2.27 \times 10^{-2}$ | $2.27 \times 10^{-2}$ |
| | AFR | $9.80 \times 10^{-3}$ | $8.53 \times 10^{-3}$ |
| *Asthma* | EUR | $1.07 \times 10^{-2}$ | $1.59 \times 10^{-2}$ |
| | AMR | $9.79 \times 10^{-3}$ | $1.47 \times 10^{-2}$ |
| | AFR | $2.53 \times 10^{-3}$ | $2.97 \times 10^{-3}$ |
| *Colorectal Cancer* | EUR | $2.86 \times 10^{-3}$ | $1.01 \times 10^{-3}$ |
| | AMR | $1.01 \times 10^{-3}$ | $1.29 \times 10^{-3}$ |
| | AFR | $8.59 \times 10^{-4}$ | $1.70 \times 10^{-3}$ |
| *Breast Cancer* | EUR | $1.56 \times 10^{-2}$ | $8.77 \times 10^{-3}$ |
| | AMR | $1.43 \times 10^{-2}$ | $6.13 \times 10^{-3}$ |
| | AFR | $3.62 \times 10^{-3}$ | $2.22 \times 10^{-3}$ |

The $R^2$ columns report both $R^2$ and pseudo-$R^2$.

**Supplementary Table 13. PGS Catalog traits evaluated in this study.**

| *Traits* | Publication (training) | # Variants | Target population | Scoring method |
|---|---|---|---|---|
| *Height* PGS002802 | Yengo, 2022 (GIANT) | 1,103,042 | EUR | SBayesC |
| | | 1,103,042 | AFR | |
| | | 1,103,042 | AMR | |
| | | 1,103,042 | EAS | |
| | | 1,103,042 | SAS | |
| *Height* | Gunn, 2024 (MVP) | 1,286,612 | EUR PGS005000 | LDpred |
| | | 1,286,612 | AFR PGS004998 | |
| | | 1,286,612 | AMR PGS004999 | |
| *HDL* PGS003773 | Zhang, 2023 (GLGC) | 1,102,319 | AFR | CTSLEB |
| *HDL* PGS002781 | Kanoni, 2022 (GLGC) | 1,239,184 | EUR, AFR, EAS, SAS | PRS-CS |
| *T2D* PGS005242 | Guo, 2025 | 1,257,036 | EUR, AFR, AMR, ASN | PRS-CSx |
| *Breast cancer* PGS000004 | Mavaddat, 2019 (BCAC) | 313 | EUR, AFR, AMR, EAS, ASN | Hard-Thresholding Stepwise Forward Regression |

PGS Catalog IDs are shown under the corresponding traits or target populations. GIANT: Genetic Investigation of Anthropometric Traits. MVP: Million Veteran Program. GLGC: Global Lipids Genetics Consortium. BCAC: Breast Cancer Association Consortium.

**Supplementary Table 18. PGS computation time of a single trait using C+T and PRS-CS in *All of Us*.**

| *Job* | CPU hours | | | | | | Ran in *AoU* platform? |
|---|---|---|---|---|---|---|---|
| | *C+T* | | *PRS-CS* (HM3 LD panel) | | *PRS-CS* (Full-scale LD panel) | | |
| | *Array* | *WGS* | *Array* | *WGS* | *Array* | *WGS* | |
| ▪ *Infer* $\hat{\beta}_{posterior}$ | N/A | N/A | 81 | 175.5 | 306 | 5,616 | No[a] |
| ▪ *Clumping* | <27 | <54 | N/A | N/A | N/A | N/A | Yes |
| ▪ *Compute PGS* | | | 27 | 80 | 11[b] | 24[b] | Yes |

a. To infer posterior effect sizes using PRS-CS, we provided summary statistics and a BIM file for each trait as input. Since no individual-level data were involved, this job was run locally outside of *All of Us* to reduce computational costs.

b. Once the panel was created, the PGS computation time using the full-scale LD panel upstream is shorter compared to the runtime with the default LD panel. This improvement is due to modifications made to the PRS-CS source code to recognize hg38 variant IDs in a format optimized for Hail genomics analysis in *All of Us*. The default PRS-CS, in contrast, only recognizes hg19 rsIDs.
